# Closed-Loop Connectivity Best Supports Angular Tuning and Sleep Dynamics in a Biophysical Thalamocortical Circuit Model

**DOI:** 10.1101/2025.08.18.670921

**Authors:** Joao VS Moreira, Fernando S Borges, Zoe Atherton, Shane R Crandall, Carmen Varela, Salvador Dura-Bernal

## Abstract

Despite recent advancements in mapping thalamic and cortical projections, the specific organization of intrathalamic and corticothalamic connectivity remains elusive. Current experimental approaches cannot definitively determine whether these connections are arranged in reciprocal (closed-) or non-reciprocal (open-loop) circuits. We developed a biophysically detailed multi-compartmental model of the mouse whisker pathway, built on anatomical and physiological data. We showed that closed-loop intrathalamic projections between the thalamocortical (TC) relay neurons in the ventral posteromedial nucleus and the inhibitory neurons in the thalamic reticular nucleus (TRN) best reproduce thalamic spiking and local field potential responses across awake and sleep states. Increasing the percentage of closed-loop projections regulates the angular tuning in the awake state, while also supporting spindle oscillations during sleep. We also showed that direct activation of closed-loop corticothalamic feedback (CT→TC and CT→TRN), simulating TC inputs, sharpens the angular tuning in the thalamus. These results contribute to resolving a long-standing question regarding the organization of intrathalamic projections, offering insights into how thalamo-cortical circuits balance precise sensory tuning with robust oscillatory rhythms across behavioral states. Moreover, all model resources are open-source and available to other researchers interested in studying thalamocortical circuits.

**Author summary:** A long standing question in the study of thalamocortical interactions is whether neurons in the thalamus form so-called open- or closed-loops when they project within the thalamus and to the cortex. In this study, we used a detailed computational model of the thalamic neurons in the whisker pathway of the mouse with realistic biophysics to investigate this question. We evaluated the impact of different connectivity arrangements in reproducing the activity of thalamic neurons observed during wakefulness and sleep. Our results show that a closed-loop circuit arrangement provides the best alternative to reproduce wake and sleep neuronal responses, highlighting the importance of computational modeling as a tool to disentangle thalamic circuit organization. We also showed that closed-loop projections from the cortex to the thalamus help further amplify the selectivity of thalamic neurons, suggesting a similar organization of corticothalamic projections. We hope our predictions can inform future experiments, and elucidate principles of thalamocortical connectivity that can be generalized across other thalamocortical motifs besides the whisker pathway and across species.

## Introduction

Understanding thalamo-cortical interactions is essential for deciphering brain function across wake and sleep states. During wakefulness, these interactions support a range of functions, including sensory relay [1,2], attentional modulation [3,4], and motor coordination [5]. During sleep, they give rise to slow oscillations and rhythmic activity such as spindles [6], particularly during stage N2 of non-rapid eye movement (NREM) sleep [7–9]. Sleep spindles play an essential role in processes like memory consolidation [10,11] and synaptic plasticity [12,13], highlighting the central role of thalamo-cortical circuits in both cognitive and physiological functions.

Despite advances in mapping thalamo-cortical projections [14–18], the architecture of intrathalamic connectivity, particularly between the excitatory thalamocortical (TC) neurons and the inhibitory neurons in the thalamic reticular nucleus (TRN), remains unclear. Current experimental methods cannot definitively resolve whether TC neurons receive direct feedback from the same TRN neurons that they innervate, forming reciprocal (closed-loop) circuits, or whether TRN feedback targets neighboring TC neurons, in a non-reciprocal (open-loop) fashion [19]. Clarifying this distinction is key to understanding how the thalamus switches between precise sensory relay during wakefulness and robust oscillatory rhythms during sleep.

The rodent whisker pathway provides an ideal system to investigate this question due to its well-defined somatotopic organization [20]. Each whisker is represented by distinct and clearly defined groups of cells from the brainstem to the cortex [21,22], which preserve a preference for projecting within this pathway, including the brainstem, thalamus, and cortex [23,24]. The ventral posteromedial nucleus (VPM) is the core whisker division of the somatosensory thalamus and is organized into somatotopic units known as *barreloids*. These barreloids receive excitatory whisker-related input from the principal trigeminal nucleus in the brainstem, which is also somatotopically arranged into corresponding units called *barrelettes* [23,25]. The receptive field of an individual TC neuron is dominated by a single “principal” whisker that evokes the strongest response. TC neurons are excitatory and send axonal projections to the inhibitory TRN and its corresponding *barrel* in layer 4 of primary somatosensory cortex (S1) [22,26].

Another key feature of the whisker pathway is the topological organization of its axonal projections, encoding whisker deflection direction selectivity as a linear map along the longitudinal axis of the thalamic barreloid [27]. This organization arises from brainstem axons that selectively innervate small clusters of neighboring TC neurons, effectively transferring the angular tuning from brainstem barrelettes to thalamic barreloids [28]. As a result, directional tuning is preserved and observable across the brainstem [1,2], thalamus [1,2,27], and cortex [29–31].

In addition, the intrinsic biophysics of TC and TRN neurons shift markedly between wakefulness and sleep. In the awake state, thalamic neurons remain relatively depolarized and fire in tonic mode, driven by Na⁺ and K⁺ currents [32]. In contrast, during NREM sleep, membrane hyperpolarization de-inactivates T-type Ca²⁺ channels [33–35], enabling low-threshold calcium spikes [36,37] and burst firing, which are brief and high-frequency spike clusters [32]. This sleep-induced hyperpolarization is driven by changes in neuromodulatory inputs, such as acetylcholine and norepinephrine [38,39], that alter ionic conductances, particularly in K⁺ channels [40]. The transition from wake to sleep is also marked by changes in synaptic strength, driven by shifts in neurotransmitter concentrations, including a two-fold increase in extracellular GABA [9,41,42].

The neurons in the thalamus are also influenced by excitatory feedback from the cortex. TC and TRN neurons receive projections from corticothalamic (CT) neurons in layer 6A (L6A) of S1 [43], exerting a rate-dependent modulation in the thalamus [44]. During wakefulness, CT neurons in L6A can be directly activated by TC projections [45–47], enabling a direct feedback mechanism to TC inputs. During sleep, these thalamo-cortical interactions are replaced by oscillatory activity, such as “Up/Down” states. These Up/Down states occur during the N2 stage of NREM sleep [48–50] and are responsible for entraining spindle activity in the thalamus, which are 8–16 Hz waxing-waning oscillations lasting 500–3000 ms [48,51].

With this in mind, we developed a biophysically detailed, multi-compartmental model of the thalamo-cortical circuit in the whisker pathway, inspired by a model of the non-barreloid thalamus [52], to investigate how intrathalamic connectivity shapes thalamic function during wakefulness and sleep. Our open-source model incorporates realistic neuronal morphologies, short-term synaptic dynamics, and anatomically derived topological connectivity. By systematically varying the connectivity between TC and TRN neurons, we show that closed-loop connectivity between TC and TRN neurons provides the best configuration to reproduce key experimental observations: precise angular tuning during wakefulness and robust spindle oscillations during sleep. Additionally, we show that closed-loop CT feedback further enhances angular tuning, highlighting the role of closed-loop projections in modulating afferent sensory inputs.

These findings help resolve a long-standing debate over the synaptic architecture of intrathalamic circuits. Moreover, it provides a framework for understanding how thalamo-cortical circuits balance sensory precision and oscillatory rhythms across behavioral states, which are principles that may extend to other sensory modalities and cortical areas.

## Results

In this work, we developed a biophysically detailed circuit model of the thalamo-cortical whisker pathway, integrating multi-compartmental morphological reconstructions, biophysical membrane properties, synaptic short-term dynamics, gap junction coupling, and topological connectivity derived from anatomical and physiological data (Fig 1). The model reproduces key features of the whisker pathway in both awake and sleep states, at the level of neuronal membrane voltages, firing rates, and local field potentials (Fig 2). The model also includes an algorithm to generate realistic brainstem inputs encoding whisker deflections (Fig 3), enabling the reproduction of thalamic angular tuning response during wakefulness (Figs 4 and 5). In addition, the model also reproduces thalamic spindle oscillations during sleep in response to CT Up/Down modulation (Fig 6).

**Fig 1.**
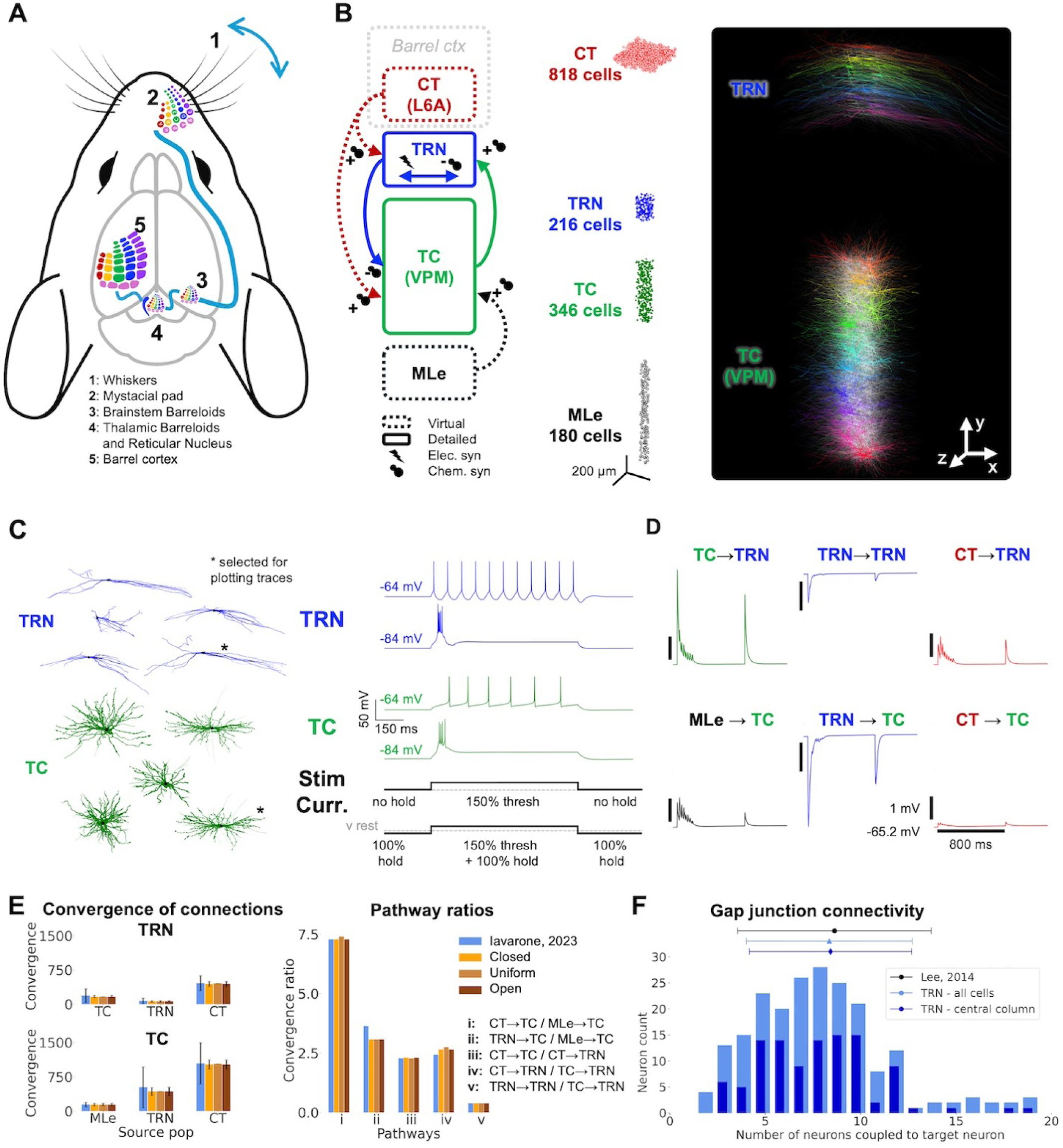
Architecture of the model of the mouse thalamo-cortical whisker pathway. (A) Schematic representation of the mouse whisker pathway. (B) Illustration of the thalamic nuclei modeled (TC neurons in a VPM barreloid and corresponding TRN region), spatial arrangement of neurons, and detailed morphological reconstructions of TC and TRN neurons. The model consists of 1,560 neurons with approximately 690k synapses across 95k cell-to-cell connections, of which 3.4k are gap junctions between TRN neurons. (C) Morphological templates and biophysical properties of modeled neurons, adapted from a computational model of the mouse non-barreloid thalamus and adjusted based on experimental data. (D) Properties and classification of chemical synapses, including pathway-specific short-term plasticity dynamics reflecting driver and modulator characteristics. Postsynaptic TC and TRN neurons were stimulated using the parameters of the 6 different synaptic models corresponding to each connection type shown in (B) [52]. The protocol consisted of 8 post-synaptic potential (PSP) stimulations every 25 ms, followed by a single PSP after 600 ms. (E) Connectivity patterns of chemical synapses within the modeled circuit. *Left:* convergence of connections received by the TC and TRN neurons. *Right:* ratio of connections from different pathways received by TC and TRN populations. Data is compared against another thalamocortical model [52]. (F) Gap junction (electrical) coupling within the TRN population. Number of neurons coupled to each TRN neuron in the model and comparison with experimental data [53]. See *Methods* for details. TC: thalamocortical; VPM: ventral posteromedial nucleus of the thalamus; TRN: thalamic reticular nucleus.

**Fig 2.**
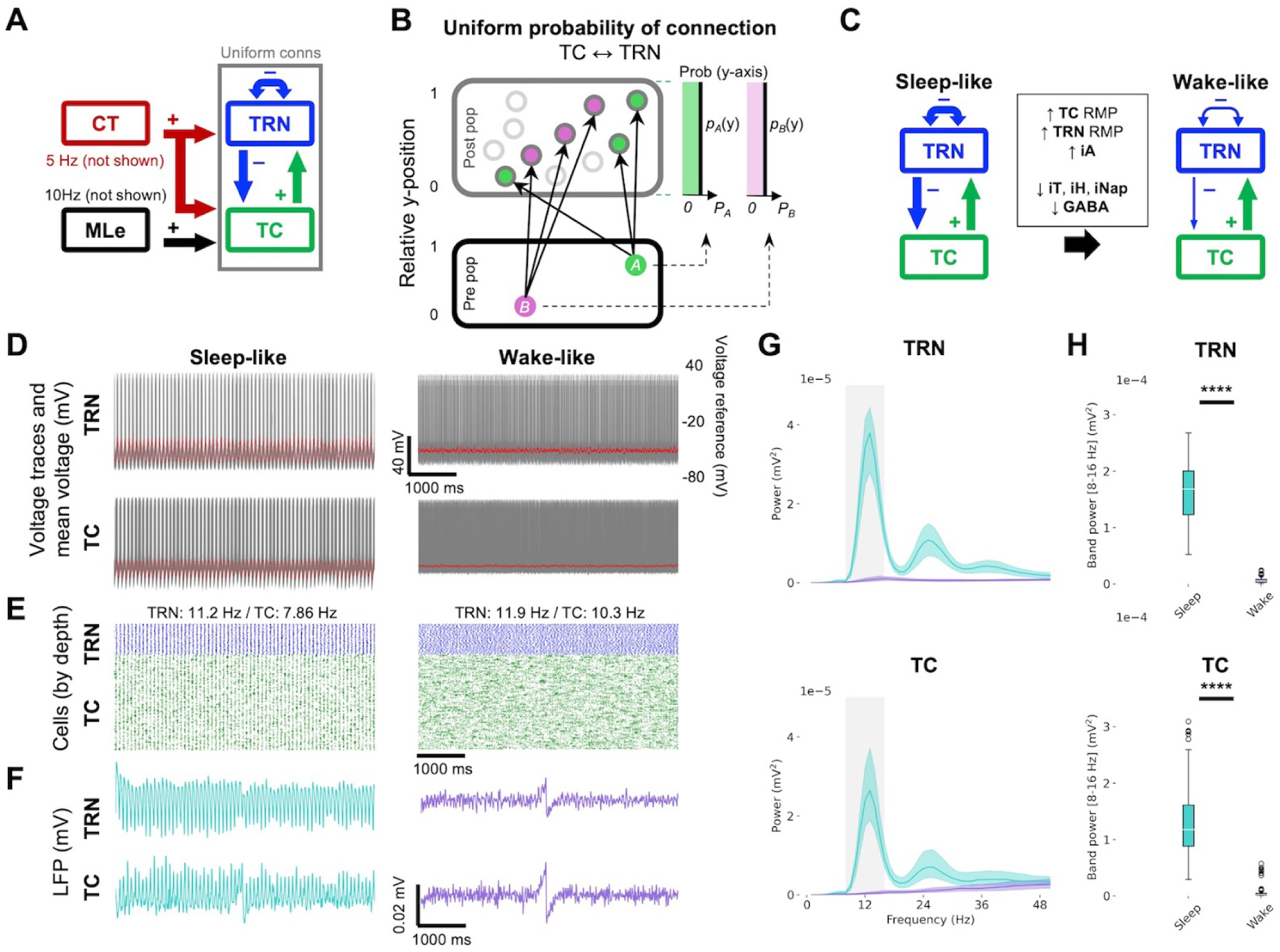
Reproduction of sleep- and wake-like dynamics in the thalamo-cortical model. (A) Circuit diagram illustrating the thalamo-cortical model architecture used to simulate baseline dynamics. (B) Diagram of uniform intrathalamic connectivity based on cell convergence data. (C) Illustration of the parameter adjustments required to transition from sleep to wake states, including changes in neuromodulation-related cellular properties and GABA neurotransmitter concentration (Table 1). (D-E) Model response during sleep and wake states, including (D) stacked membrane voltage of TRN neurons (grey) and average membrane voltage in TRN (red); and (E) TC and TRN spiking raster plots. (F) TC and TRN LFP recordings. (G) PSD of the TC and TRN LFP for sleep- vs wake-like conditions, with the spindle frequency range (8–16 Hz) highlighted in grey (Solid line: mean value; shaded intervals: standard deviation). (H) Statistical comparison of LFP power in the spindle frequency range (8–16 Hz) between sleep and wake states. Significant decrease in spindle power was observed in the wake state compared to the sleep state (Two-sided Mann-Whitney U test; TRN: p-value=3.92e-95****, TC: p-value=1.11e-94****). TC: thalamocortical; TRN: thalamic reticular nucleus; LFP: local field potential; PSD: power spectral density; GABA: gamma-aminobutyric acid.

**Fig 3.**
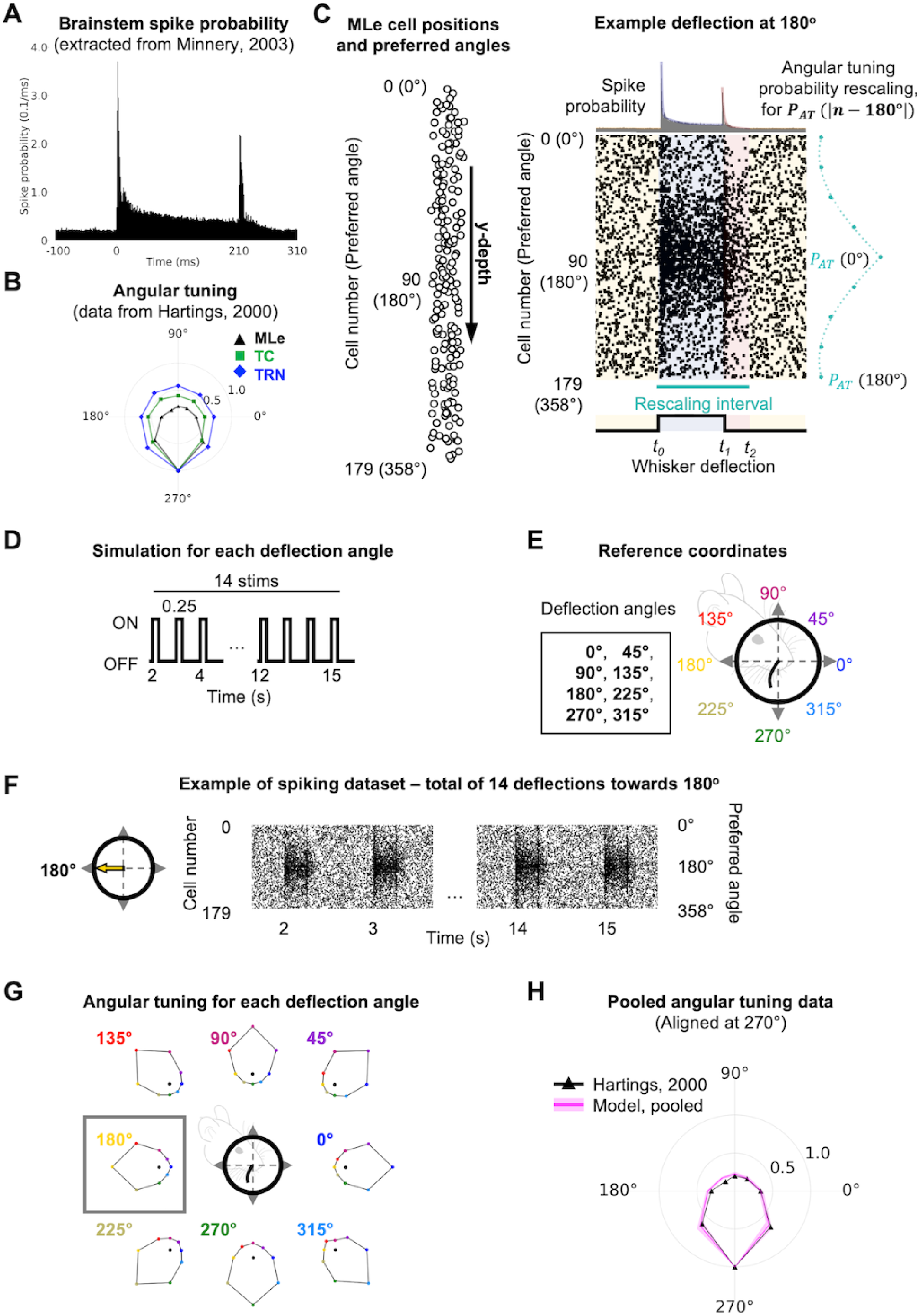
Reproducing brainstem barrelette activity and angular tuning with a realistic whisker deflection model. (A-B) Fitting of experimental brainstem activity data during whisker deflection, including (A) PSTH of spike probabilities [2] and (B) angular tuning [1]. (C) Illustration of cell positions and maximally responsive angles, and spiking activity sampled based on fitted spike probability and angular tuning models, representing 180 spike generators (2° resolution, 0°–358°). (D) Protocol for generating spiking datasets representing whisker deflections at eight angles (0° to 315° in 45° increments). (E) Reference coordinate map and deflection angles used for dataset generation. (F) Example raster plot showing generated spike activity for a 180° whisker deflection. (G) Angular tuning maps corresponding to each deflection angle depicted in (E). Grey box shows the angular tuning map for the spikes in (F). (H) Average angular tuning map derived from the whisker deflection model compared to experimental data (aligned to 270° reference angle) [1]. For detailed descriptions of spike sampling and angular tuning calculation methods, see *Methods* and *Supporting information*. PSTH: peristimulus time histogram.

**Fig 4.**
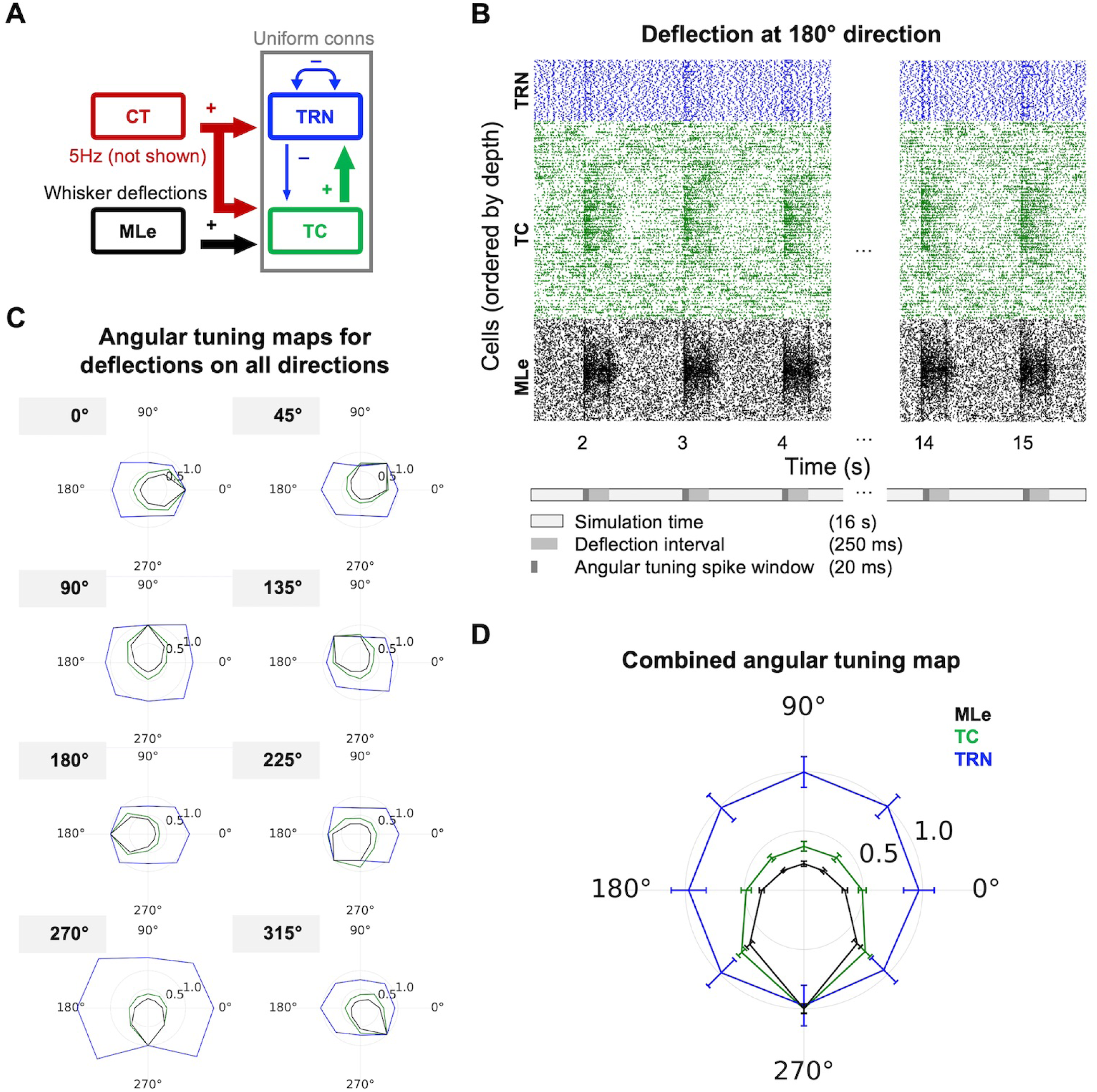
Angular tuning response in a thalamo-cortical circuit with uniform connectivity. (A) Circuit diagram illustrating the thalamo-cortical model in the awake-like state, with uniform connectivity between TC and TRN populations. (B) Raster plots of spiking activity in MLe, TC, and TRN cell populations in response to repeated whisker deflection inputs presented at 1-second intervals (CT 5 Hz baseline spiking not shown). (C) Angular tuning maps depicting responses from MLe, TC, and TRN populations across the eight whisker deflection directions. (D) Average angular tuning map computed by aligning responses from all deflection directions to a common reference (270°), pooling, and normalizing spike counts for each population. See *Methods* for additional details on spike extraction, alignment, and angular tuning calculations. MLe: medial lemniscus; TC: thalamocortical; TRN: thalamic reticular nucleus; CT: corticothalamic.

**Fig 5.**
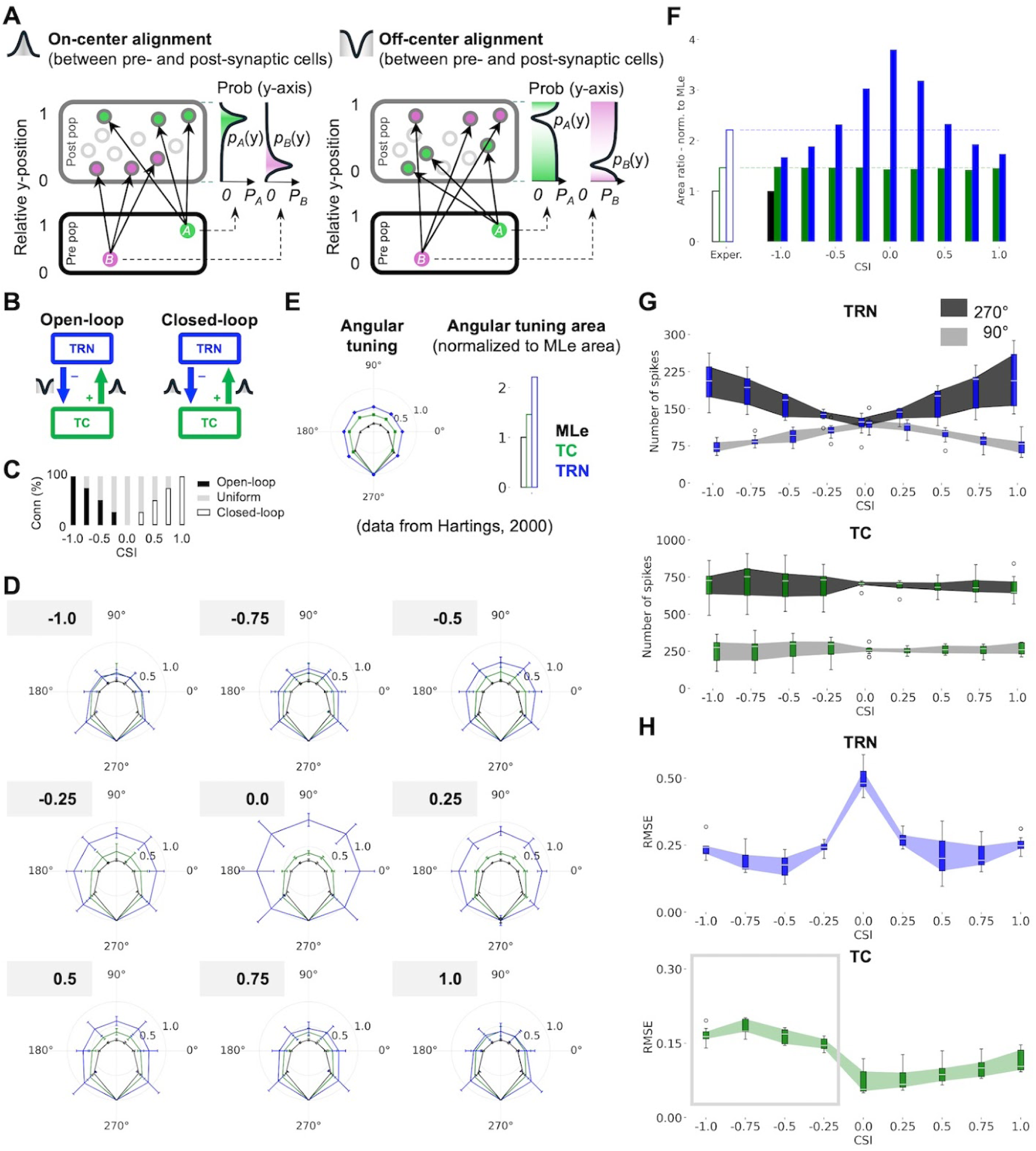
Effects of intrathalamic connectivity in TC and TRN angular tuning. (A) Schematic representation of on-center and off-center alignment of connections between pre- and post-synaptic populations. (B) Definition of open-loop and closed-loop configurations based on combinations of on- and off-center alignment of feedforward and feedback projections. (C) Diagram of intrathalamic circuit configurations tested, ranging from fully open-loop (CSI = -1.0), uniform (CSI = 0.0), to fully closed-loop (CSI = 1.0), varied in steps of 25%. (D) Angular tuning maps for each circuit configuration. Generated using methods described in Fig 4. (E) Experimental angular tuning data used to validate the model [1]. Area ratio calculated relative to MLe for TC and TRN populations. (F) Quantification of angular tuning map areas normalized to MLe tuning area, highlighting changes across connectivity conditions. (G) Spike count analysis of aligned and opposite neurons (maximally responsive to 270° and 90°, respectively) for each circuit configuration, illustrating differential spiking patterns related to connectivity. Boxplot and shading reflect variability in the total number of spikes of aligned and opposite neurons across different deflection directions (aligned to 270°), for each CSI. (H) AT_RMSE_ of TC and TRN populations across all circuit configurations. Boxplot and shading reflect variability across all angles (except for 270°, which is the normalization reference) for each CSI. Statistical analyses and details are provided in Table S1 of *Supporting Information*. MLe: medial lemniscus; TC: thalamocortical; TRN: thalamic reticular nucleus; CT: corticothalamic; CSI: connectivity symmetry index; AT_RMSE_: root mean squared error of the angular tuning.

**Fig 6.**
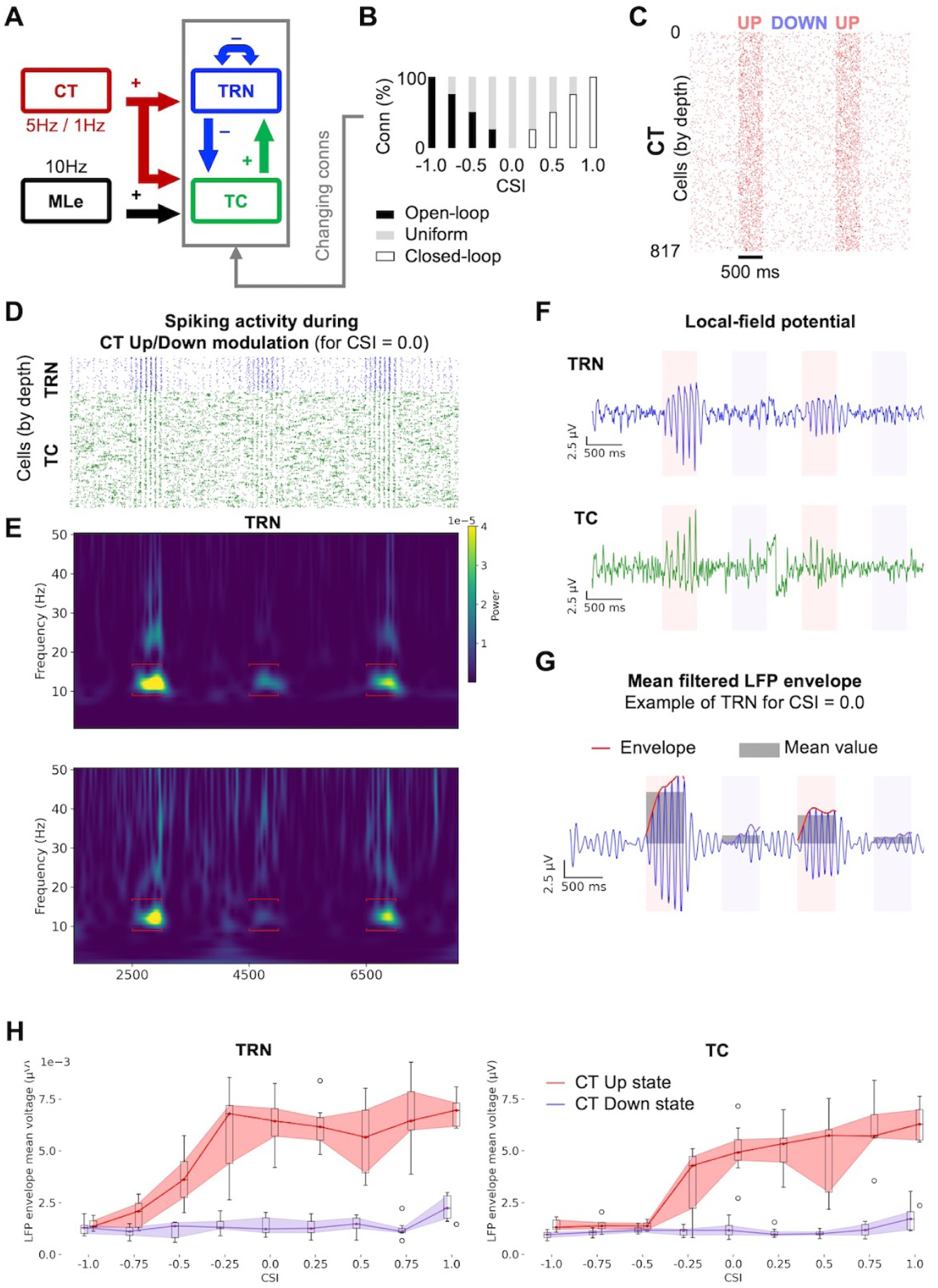
Open-loop connectivity disrupts spindle oscillations during CT Up states. (A) Circuit diagram illustrating the model configured for the sleep state. (B) Intrathalamic circuit configurations tested, combining open-loop, uniform, and closed-loop projections. (C) Spiking raster plot showing CT Up/Down states, with cells spiking at 5 Hz and 1 Hz, respectively. (D) Spiking raster plots of TC and TRN populations during CT Up/Down modulation. See Fig. S1 for a visual comparison across all circuit configurations. (E) Spectrograms highlighting the thalamic population activity, with red brackets indicating the spindle frequency range (8–16 Hz). (F) LFP signals from TC and TRN populations, and segmentation of Up/Down states. The central part of the Down segment was also segmented to ensure that any oscillations returned to baseline after the Up state. (G) Segment of the LFP signal filtered within the spindle frequency range. Up and Down epochs were used to calculate the signal envelope, and the mean envelope amplitudes were computed to quantify spindle oscillations. (H) Distribution of mean LFP envelope amplitudes during CT Up and Down states across all intrathalamic circuit configurations (CSI: [-1.0, 1.0]), demonstrating the differential impact of closed- vs open-loop connectivity on spindle oscillations. See Table S2 in *Supporting information* for detailed statistical analyses. TC: thalamocortical; TRN: thalamic reticular nucleus; CT: corticothalamic; CSI: connectivity symmetry index; LFP: local field potential.

**Table 1:** Model parameters used to simulate the sleep vs wake states.

| ***i* ) Simulation parameter** |  | **Sleep** | **Wake** |
| --- | --- | --- | --- |
| **Rescale maximum conductance** | *iT* | 1.0 | 0.25 |
|  | *iH* | 1.0 | 0.25 |
|  | *iNap* | 1.0 | 0.25 |
|  | *iA* | 1.0 | 1.5 |
| **RMP (mV)** | *TC* | -61 | -58 |
|  | *TRN* | -71 | -58 |
| **Rescale GABA strength** | *TRN→TC* | 1.0 | 0.3 |
|  | *TRN→TRN* | 1.0 | 0.5 |
| **Rescale AMPA/NMDA strength** | *MLe→TC* | 3.0 | 3.0 |
|  | *TC→TRN* | 0.5 | 0.5 |
|  | *CT→TC* | 1.0 | 1.0 |
|  | *CT→TRN* | 1.5 | 1.5 |
| ***ii* ) Tuning targets** |  | **Sleep** | **Wake** |
| **Firing Rate (Hz)** | *TC* | 6–9 | 9–15 |
|  | *TRN* | 9–15 | 9–15 |
| **Spindle Frequency (Hz)** | *TC* | 8–16 | - |
|  | *TRN* | 8–16 | - |
| ***iii* ) Model's response** |  | **Sleep** | **Wake** |
| **Firing Rate (Hz)** | *TC* | 7.86 | 10.3 |
|  | *TRN* | 11.2 | 11.9 |
| **Spindle Frequency (Hz)** | *TC* | 13 (peak) | - |
|  | *TRN* | 13 (peak) | - |

We leveraged these features to investigate the organization of intrathalamic connections (Figs 5 and 6) and the influence of direct corticothalamic activation on angular tuning (Fig 7). This open-source model represents a significant step forward in studying thalamo-cortical dynamics, offering a robust platform to investigate how structural changes in thalamic circuitry impact sensory processing. In the following sections, we explore in detail how thalamic dynamics and function, such as angular tuning, sleep oscillations, and corticothalamic modulation, are related to thalamic circuit connectivity and what the implications are for encoding sensory information.

**Fig 7.**
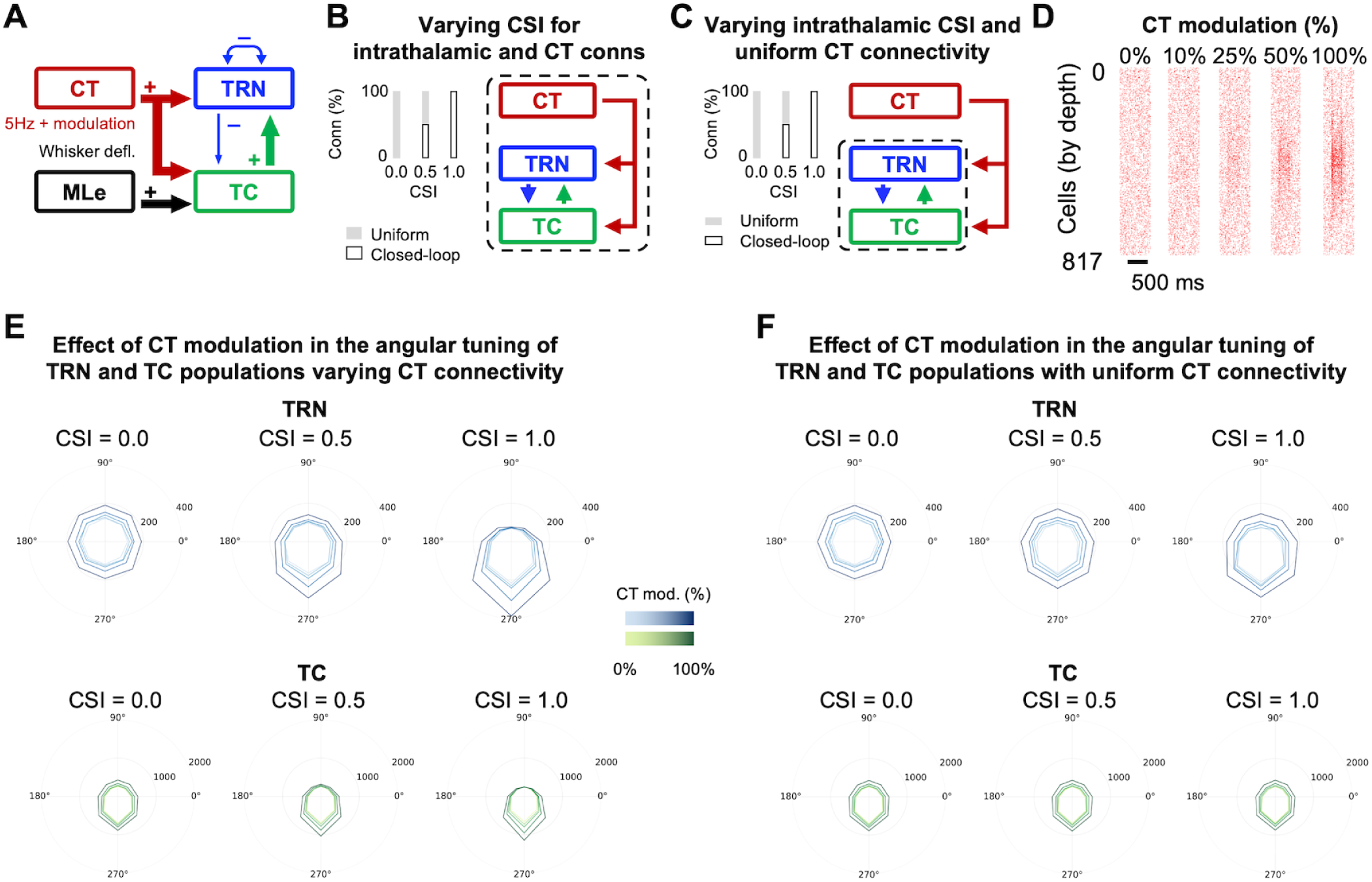
Closed-loop CT modulation selectively enhances angular tuning in thalamic circuits. (A) Schematic illustrating simulation of monosynaptic CT activation, with angular-tuned CT modulation. (B-C) Circuit configurations tested: uniform (CSI = 0.0), mixed (CSI = 0.5), and fully closed-loop (CSI = 1.0), involving the intrathalamic and corticothalamic connections (B), or the intrathalamic connections only, keeping corticothalamic connections uniform (C). (D) Illustration of different levels of CT modulation, representing proportions (0%, 10%, 25%, 50%, 100%) of total neurons showing angular-tuned responses (example of a 180° whisker deflection). (E-F) Differential analysis of spiking activity in TRN and TC neurons under increasing levels of CT modulation. Analysis of spiking activity considered aligned neurons (270°) versus opposite neurons (90°) relative to the target angle. Side-by-side comparison of the influence of CT modulation in the circuits with closed-loop intrathalamic and corticothalamic connections (E), versus the circuit with closed-loop intrathalamic connections and uniform CT connections (F). See Figure S2 and Tables S3 and S4 in *Supporting information* for detailed comparison and statistical analyses. TC: thalamocortical; TRN: thalamic reticular nucleus; CT: corticothalamic; CSI: connectivity symmetry index.

### Detailed computational model of the thalamo-cortical circuit in the mouse whisker pathway

The computational model of the whisker pathway circuits developed in this study includes 1,560 neurons, including 562 detailed morphological reconstructions of TC and TRN neurons within a thalamic barreloid in VPM and its projecting region in TRN (Fig 1A-B). The morphological and biophysical properties of these neurons were extracted from a computational model of the mouse non-barreloid thalamus [52,54] (Fig 1C) and adapted to represent the barreloid structure and TRN sector based on experimental data [2,21,27,55–58]. The remaining 998 neurons are implemented as artificial generators of precisely-timed spiking patterns and divided between the brainstem (180) and cortical (818) populations.

These neurons communicate via chemical synapses with short-term plasticity reflecting pathway-specific properties, such as driver and modulator characteristics [58], and via dendro-dendritic electrical synapses in the TRN population [59] (Fig 1D). These pathway-specific properties are used to determine the connectivity rules between each pre- and post-synaptic population. Driver synapses (MLe→TC and TC→TRN) carry the main afferent information to the postsynaptic neuron and present short-term depression. Conversely, modulator synapses influence the firing properties of the postsynaptic neuron, without being responsible for triggering the action potential, and these synapses can be either facilitating (CT→TC and CT→TRN) or depressing (TRN→TC and TRN→TRN) [60]. Driver synapses were placed in the soma and proximal dendrites of postsynaptic cells, and modulator synapses targeted the intermediate and distal zones of the dendritic tree [60,61]. In total, the circuit includes approximately 690k synapses across 95k cell-to-cell connections, of which 3.4k are gap junctions between TRN neurons (see detailed description in *Methods*). The average number of cell-to-cell connections for TC and TRN neurons is shown in detail in Fig 1E, and the distribution of intra-TRN gap junctions is shown in Fig 1F.

### Unified thalamo-cortical model reproduces sleep and wake state dynamics

To evaluate the baseline circuit dynamics, we characterized the model’s response under 10 Hz brainstem input and 5 Hz cortical feedback (Fig 2A) [52]. For the baseline characterization, we initially assumed uniform intrathalamic connectivity; the influence of different circuit connectivity configurations on the model’s response is explored in later sections. Hence, the probability of connection between neurons in the pre- and postsynaptic populations was the same, regardless of their spatial distribution. The number of connections received by each cell was determined based on cell convergence data for the TC→TRN and TRN→TC pathways [52] (Fig 2B).

By default, the model exhibits synchronized activation of TC and TRN populations in the spindle frequency range (8–16 Hz) [48], which are consistent with a sleep-like state. These properties were inherited from the original implementation of the model, which focused on the non-barreloid sensory thalamus [52]. This result suggests that the convergence and divergence matching in our implementation (Fig 1E) preserves the network balance, yielding similar activity in the sleep-like state.

Following, we simulated a wake-like state in the model state by adjusting the resting membrane potential (RMP) of the TC and TRN neurons, and scaling the strength of key synaptic pathways and GABAergic input to reduce oscillatory activity in the circuit (Fig 2C). These modifications are grounded in previous experimental and modeling results [9,38,40–42,62,63], and reflect a phenomenological representation of the wake-like activity, with abolishment of oscillatory activity and transition to a linear regime. Most notably, we rescaled the conductance of the T-type Ca^2+^ current (iT) and the hyperpolarization-activated cation current (iH), which are the key players in generating sustained oscillatory activity in the intrathalamic networks [63], and the persistent sodium current (iNap), to prevent runaway excitability following the positive RMP shift in TC and TRN [64]. We also increased the conductance of the A-type calcium current (iA), which is known to act as a “shock-absorbing” current, reducing network synchronization, which can trigger oscillatory activity [65]. Moreover, we rescaled the strength of GABAergic inputs within the TRN and to the TC neurons, based on previous reports that measured changes in its concentration during sleep and awake states [41]. See Table 1 for a detailed summary of the cellular and synaptic properties that reflected the change from sleep-like to wake-like state. These manipulations abolished oscillatory activity and synchronized firing (Fig 2D-F), as evidenced by the reduction in spindle power (Fig 2G-H). These two states form the foundation for subsequent experiments, in which we investigate how intrathalamic connectivity shapes circuit dynamics.

### Whisker deflection model accurately captures brainstem barrelette activity and angular tuning

To produce realistic thalamic activity, it is essential to drive the circuit model with biologically realistic brainstem inputs. To achieve this, we developed an empirical spiking model of a brainstem barrelette (hereafter referred to as the “whisker deflection” model), guided by experimental data on peristimulus time-histogram (PSTH) responses to whisker deflections (Fig 3A) [2] and angular tuning (Fig 3B) [1] of these neurons. The whisker deflection model comprises 180 artificial spike generators, each maximally responsive to a 2° sector in polar coordinates determined by their depth, consistent with experimental estimates [27,66]. The model can represent whisker deflections in any direction by scaling the spiking probability of each neuron based on its relative position along the y-axis (Fig 3C).

To validate the whisker deflection model, we produced eight 16-second stimulation datasets (0° to 315°, in 45° increments) (Fig 3D-F), computed their angular tuning responses (Fig 3G), and compared the average tuning response to experimental data (Fig 3H) [1]. During baseline activity, the background inputs provided a uniform 20Hz stimulation [52], with a transient increase in firing during each whisker deflection, totaling ∼30Hz altogether. These firing rates were based on PSTH experimental recordings for baseline and whisker deflection spike probabilities [1]. As for the TC and TRN neurons, the firing rate, including whisker deflections, was around 13 Hz for both populations, also approximating experimental findings [1]. The result is a realistic computational model of brainstem inputs that provides the thalamo-cortical model with accurate spiking activity representing whisker deflections in eight different directions (Fig 3H). See *Methods* and *Supporting information* for further details on the whisker deflection algorithm, spike probabilities, and the calculation of angular tuning. Additional information on generating these spike times can be found in the source code linked with this publication.

### Thalamo-cortical model exhibits angular tuning in the awake state

To characterize the circuit’s response to whisker deflection inputs in the awake state, we measured the average angular tuning in the thalamus. This process is illustrated in Fig 4, considering a uniform connectivity between thalamic populations as a starting point (Fig 4A). Experimental studies consider the first 20 ms window of spiking to characterize the angular tuning in response to a whisker deflection, which captures the transient increase in spiking activity in thalamus and brainstem (see example in Fig 3A). Therefore, for each of the eight whisker deflection directions, we extracted the first 20 ms of spiking activity across 14 deflection events. We aligned all responses to a common reference direction (270°), pooled the data across cells, and normalized it by the total spiking activity at 270° for the MLe, TC, and TRN populations [1,2] (Fig 4B-C). The result was a combined angular tuning map, representing the average tuning of each population in the model (Fig 4D).

The angular tuning map shows a clear transference of the angular tuning from the MLe (brainstem) to the TC (thalamic relay) population, driven by the topological connectivity of these projections. The TRN population, however, shows no preferential tuning, which is expected, since both the feedforward TC→TRN and the feedback TRN→TC connections are uniformly distributed. In the following section we will explore the effects of gradually replacing these uniform connections by either open- or closed-loops, and its influence in the angular tuning.

### Closed-loop intrathalamic connectivity best supports angular tuning in the thalamus

Although the spatial distribution of brainstem barrelette projections to thalamic barreloids is well characterized [27,28,67], the pattern of neuron-to-neuron connectivity within the thalamus remains elusive. Specifically, a key open question is whether feedforward and feedback projections between TC neurons across different thalamic relay nuclei (e.g., VPM barreloids) and the TRN follow a closed-loop or open-loop organization [14–19] (Fig 5A-B). In a fully open-loop circuit, the intrathalamic feedforward connections (from TC→TRN) are on-center (targeting the same region/cells where it received input from) (Fig 5A), and the intrathalamic feedback connections (from TRN→TC) are off-center (targeting outside of the input source) (Fig 5A). Conversely, in a fully closed-loop circuit, both the intrathalamic feedforward and feedback connections are on-center.

To investigate intrathalamic synaptic organization, we systematically replaced the uniform connections between TC and TRN populations by either open- or closed-loop connections, and assessed their effects on angular tuning in the awake state (Fig 5C). To do this, we introduced a connectivity symmetry index, which is defined as the signed fraction of uniform TC↔TRN connections that were replaced by either open-or closed-loop connections:

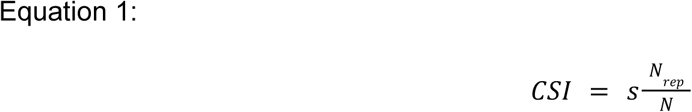

Where:

*N* is the total number of initially uniform connections, *N*_*rep*_ is the number of connections replaced, and *s* = 1 for replacement by closed-loop connections, and *s* =− 1 for replacement by open-loop connections. Thus, CSI = 0.0 denotes fully uniform connectivity, CSI = 1.0 denotes complete replacement by closed-loop connectivity, and CSI = -1.0 denotes complete replacement by open-loop connectivity. We simulated circuits with CSIs ranging from -1.0 to 1.0 in increments of 0.25 (equivalent to replacing 25% of connections), measured how connectivity affects angular tuning (Fig 5D), and evaluated it against experimental data (Fig 5E).

To compare the differences in angular tuning across different connectivity arrangements, we calculated the “area ratio” for the angular tuning maps, dividing the area of the TC and TRN angular tuning maps by the area of the MLe angular tuning map. We found that the TC area ratio remained unchanged across circuit manipulations (Fig 5F), due to the dominance of topological MLe→TC feedforward projections, implemented based on detailed anatomical studies [28,67]. In contrast, TRN neurons showed a decrease in the area ratio as uniform connections were replaced by either open- or closed-loop, indicating a sharpening in the tuning (Fig 5F). This effect was attributed to identical feedforward TC→TRN on-center connectivity, which is shared across all opposing CSI configurations (e.g., CSI = -0.5 and CSI = 0.5 share the same intrathalamic feedforward connectivity, but opposite feedback connections).

To evaluate whether the changes in the angular tuning maps reflected an increase in tuning, we compared spike counts for neurons most responsive to the deflection angle of interest (remapped to 270°, termed “aligned”) with those most responsive to the opposite angle (remapped to 90°, termed “opposite”) (Fig 5G). In TRN, increasing feedforward on-center connectivity boosted firing rates of aligned neurons and reduced activity of opposite neurons (Fig 5G; Table S1i), regardless of whether feedback was off-center (open-loop) or on-center (closed-loop). In TC, aligned neurons consistently fired more under every condition, reinforcing the role of MLe inputs in driving angular tuning in this population (Fig 5G; Table S1i).

Moreover, the spreading in the number of TC neuron spikes in open-loop circuits (CSI < 0) suggests a higher variability in the spiking activity compared with closed-loop circuits (CSI > 0) (Fig 5G). To evaluate that, we assessed angular tuning variability by calculating the root mean square error between the model and experimental angular tuning maps (AT_RMSE_). Our results show that TC neurons in open-loop circuits (CSI < 0) showed higher AT_RMSE_ relative to both uniform and closed-loop circuits (CSI ≥ 0) (Fig 5H, box; Table S1ii-iii). Besides that, TRN neurons in both open- and closed-loop circuits exhibited lower AT_RMSE_ compared to the uniform circuit (Fig 5H, top; Table S1ii).

In summary, we show that closed-loop connections can successfully transfer the TC→TRN tuning without significantly increasing the variability in the spiking activity. While a uniform circuit also keeps variability low, it fails to transfer the TC→TRN tuning. Taken together, these results highlight the strong influence of feedforward on-center input from TC, and suggest that off-center TRN feedback is detrimental for TC angular tuning. It also shows that we can use the angular tuning response as a metric to assess the level of feedforward topological connectivity in the circuit. In the following section, we extend our analysis of circuit connectivity to the sleep state, examining the role of TRN feedback in generating spindle oscillations.

### Closed-loop intrathalamic connectivity supports spindle oscillations under CT Up/Down modulation

A hallmark feature of thalamic circuits during sleep is their ability to engage spindle oscillations in response to cortical Up/Down modulation. To assess how intrathalamic connectivity influences this phenomenon, we tested the response of different circuit configurations in the sleep-like state (Fig 6A-B) to CT Up/Down modulation (500/1500 ms, respectively) (Fig 6C).

In Fig 6D-E, we show the spiking activity and spectral characteristics of a uniform thalamic network (CSI = 0.0), highlighting the increased power within the spindle frequency range during CT Up states. To systematically evaluate the effect of connectivity on the generation of sleep spindles, we quantified spindle power by band-pass filtering (8–16 Hz) the local field potential (LFP) signals from the TC and TRN populations. The filtered LFP was then subdivided into Up (500 ms) and Down (500 ms) epochs, and we measured the mean LFP envelope amplitude (Fig 6F-G, see legend for details). We used this measure to compare the spindle-range LFP amplitude in Up and Down states across different circuit connectivity configurations (Fig 6H) (See Fig. S1 for a visual comparison of the spiking activity across all circuit configurations).

Our findings indicate that closed-loop connectivity best supports spindle oscillations in the barreloid thalamo-cortical circuit, while increasing open-loop connectivity disrupts them. Specifically, open-loop projections (CSI < -0.25) weaken spindle power in Up states relative to uniform connectivity (CSI = 0.0), for both TC and TRN, with the effect becoming more pronounced as open-loop connectivity increases (Fig 6H; Table S2i). By contrast, TC and TRN closed-loop circuits exhibit stronger Up state spindle power than an equivalent open-loop circuit, as uniform connections are gradually replaced (Fig 6H; Table S2ii). Moreover, increasing the number of open-loop connections reduces the difference in spindle-range LFP amplitude between Up and Down states, but the opposite is not true for closed-loop connections (Fig 6H, Table S2iii).

In summary, these results highlight that closed-loop connectivity supports network synchronization, working as a viable mechanism for sustaining spindle oscillations. Conversely, open-loop circuits promote propagation of activity, supporting a traveling wave-like type of activity (Fig S1).

### Closed-loop fast CT feedback sharpens angular tuning

Previous experimental work has shown that TC inputs can monosynaptically activate CT neurons, in turn providing feedback to TC and TRN neurons [45–47,68]. However, the functional significance of this fast feedback remains poorly explored. In the barrel system, such fast feedback pathways may form a positive loop that amplifies topologically organized signals originating from downstream brainstem inputs. To test this idea, we simulated the fast activation of CT neurons by adding a delayed (13.1 ms) [45] angular-tuned response to the CT spiking population (Fig 7A). We evaluated the effect of this fast activation on angular tuning across three circuit configurations, varying the connectivity of the TC↔TRN circuit, and the CT→TC and CT→TRN feedbacks from uniform to fully closed-loop (CSI = 0.0, CSI = 0.5, and CSI = 1.0) (Fig 7B). We also evaluated the case where only the TC↔TRN circuit connectivity varies, while the CT→TC and CT→TRN feedbacks remain fully uniform (Fig 7C), like in previous simulations (Figs 5-6).

In both cases (Fig 7B-C), we modulated the response of the CT neurons by adding an artificial angular tuning feedback representing the activation of CT afferents. For each circuit configuration, we ran different instances of the model progressively increasing the fraction of activated CT neurons (0%, 10%, 25%, 50%, 100%) (Fig 7D), to determine how selective recruitment of CT feedback influences angular tuning.

To quantify the effect of angular-tuned CT feedback in the thalamic network, we compared the spiking activity of neurons *aligned* and *opposite* to the deflection angle (for visualization purposes, we remapped all aligned responses to 270° and opposite to 90°) (Fig 7E-F). This allowed for a visual comparison of the changes in spiking by inspecting the differences in size and shape of the tuning maps for TRN and TC populations under different circuit configurations and levels of CT modulation (Fig 7E-F). Our findings show that circuits including both intrathalamic and corticothalamic closed-loop connections (CSI = 1.0) exhibited the highest differences in spiking between aligned and opposite neurons (Fig 7E), particularly for the TRN population (Fig S2A). Notably, aligned TRN neurons showed significantly higher firing rates with increasing CT modulation, while the firing rate of opposite neurons remained unchanged (Fig S2A; Table S3i). This difference in TRN spiking between aligned and opposite neurons was diminished in the circuit with mixed intrathalamic and corticothalamic closed-loop connectivity (CSI = 0.5), and not present in the uniform circuit (CSI = 0.0), underscoring the critical role of intrathalamic and corticothalamic closed-loop circuits in fine-tuning angular selectivity.

Lastly, while the spiking activity of aligned neurons increased with CT feedback across all conditions, in the closed-loop circuit (CSI = 1.0), the spiking activity of opposite neurons remained similar for both TC and TRN populations (Figs 7E,S2A; Table S3ii). In contrast, this effect was not present in the circuits with uniform CT feedback (Figs 7F,S2B; Table S4ii), where the spiking activity of opposite neurons increased with CT modulation, regardless of thalamic connectivity (CSI = 0.0, 0.5, and 1.0). This result suggests that closed-loop connectivity in the CT feedback can further sharpen the thalamic angular tuning.

Taken together, our findings reveal that angular-tuned CT feedback is a powerful mechanism for sharpening angular tuning in circuits containing closed-loop connectivity, with particularly strong effects in TRN. Moreover, this effect is more pronounced when CT connectivity is organized in a closed-loop.

## Discussion

### Key findings and contributions

In this work, we developed an experimentally validated, detailed computational model of the thalamo-cortical circuit in the mouse whisker pathway to explore the effect of intrathalamic connectivity in sensory relay during wakefulness and spindle oscillations during sleep. Our results demonstrate that intrathalamic closed-loop connectivity provides the better arrangement to support both functions. While the balance between closed-loop and uniform projections influenced the sharpness of angular tuning, both configurations promoted spindle activity. In contrast, open-loop connectivity was less effective in reproducing the experimentally observed angular tuning and sleep spindle oscillations. Additionally, fast CT feedback modulated thalamic angular tuning maps, resulting in sharpening of sensory selectivity only when closed-loop connectivity was present in both intrathalamic and cortico-thalamic circuits. Below, we discuss the key model features and results, and the broad implications of these findings for understanding the relationship between sensory processing, sleep regulation, and neural dynamics.

Our model combines several key features of thalamo-cortical circuits that were not previously available under a unified model, enabling it to investigate the effect of cellular mechanisms and connectivity on circuit dynamics and function. For example, previous models often lacked morphological details, realistic connectivity, and/or realistic brainstem and CT feedback [52,69,70]. Our model reproduces the structure of a thalamic barreloid in the VPM nucleus and corresponding TRN region, integrating morphologically detailed TC and TRN neuronal populations with accurate spatial distributions, population-specific biophysical parameters, synaptic short-term dynamics, precise spatial targeting, and realistic spiking inputs from the brainstem and cortical inputs, and intrathalamic topological connectivity. These features collectively enabled us to reproduce stimulus-locked spike timing, fast sensory processing, and angular tuning curves during wakefulness, and synchronized spindle-like oscillations during sleep states with high fidelity to experimental observations [1,2,48,52]. Importantly, the model can simulate both the active wakefulness and sleep-like responses and state-dependent oscillatory dynamics by adjusting cellular properties and synaptic weights consistent with experimental observations [9,38,40–42,62].

Our novel brainstem whisker deflection model directly innervates the thalamus, providing a framework to systematically assess how features such as intrathalamic connectivity influence sensory processing, particularly angular tuning to whisker deflection. This whisker deflection model was built based on data mapping the projections from the brainstem barrelettes to the thalamic barreloids, focusing on the transfer of whisker deflection information [27,28,56,67]. The CT circuit was dimensioned to represent the number of corticothalamic cells in a cortical infrabarrel in layer 6 of the barrel cortex, enabling a systematic exploration of the CT modulation that regulates the activity of TC and TRN neurons [68,71].

Our comprehensive model provides a platform for exploring hypotheses about thalamo-cortical dynamics and their role in sensory information processing [1,2] and sleep-related activity, providing insights that are sometimes inaccessible via experiments. For example, combining neuron morphologies and ion channel kinetics with population-level connectivity allows for an in-depth exploration of how individual neuron properties influence overall circuit behavior, such as synaptic integration [72–74] and the effects of CT and TRN localized inputs on TC neuron responsiveness [61,75]. Including short-term plasticity in excitatory and inhibitory synapses enables the model to reproduce realistic, time-dependent changes in sensory processing and the modulation of thalamo-cortical interactions over time [52,76]. This feature is particularly relevant for capturing the transient changes in communication efficiency between neurons during different states, such as wakefulness and sleep, allowing researchers to study the role of synaptic plasticity in phenomena like sensory adaptation and the emergence of sleep spindles.

### Limitations of the study

Although our model represents one of the most detailed thalamic circuit models to date, we acknowledge a set of limitations. First, our model reproduces the thalamic network associated with a single whisker, while the mouse features around 30 whiskers on each side of the body [77], and thalamic barreloids are subject to modulation by neighboring CT barrels via lateral projections in the TRN [78]. Second, the artificial CT feedback used here is unidirectional, overlooking the reciprocal and dynamic nature of cortico-thalamic loops (more on that in a following section). Third, although neuromodulatory influences are accounted for, explicit biophysical modeling of cholinergic and noradrenergic pathways was not included. Additionally, other inputs from brainstem and cortical areas were omitted, limiting the model’s functionality.

We also acknowledge that there are some methodological limitations in our experimental design. We explored a relatively limited set of connectivity arrangements based on available data, but which may not encompass all biologically plausible network configurations. Additionally, the strategy we adopted to create the open- and closed-loop networks does not take into account how axonal projections are distributed within neighboring cells. This means that neurons that are very close together in space will not be differentially targeted when receiving a feedback projection, since we do not control for single-cell pairwise open- or closed-loops. Evidence of such an organization, where a TRN neuron receives input from a TC neuron and sends feedback to its neighbor, has been previously shown [79–81].

Moreover, we did not evaluate the influence of gap junctions in combination with the intrathalamic connectivity in this study. We opted for that to keep the study concise, since the effects of gap junctions have been extensively explored in the much larger non-barreloid thalamic network that our model was based on [52].

Future studies could address these limitations by incorporating a multi-whisker configuration, bidirectional cortico-thalamic interactions, improved neuromodulatory inputs, and refinement of cell-to-cell connectivity. Experiments explicitly designed to validate model predictions might include optogenetic manipulations and paired recordings from neighboring TRN and TC cells, to selectively activate circuit motifs and test predicted pairwise connectivity. These targeted experimental validations would significantly enhance the predictive value and biological relevance of our computational framework.

Another future improvement could be to model the effects of sensory adaptation or potentiation in the somatosensory thalamus, and how it shapes the angular tuning during repeated whisker deflections. This process involves longer timescales, and it has been linked to the recruitment of g-protein-coupled receptors in the somatosensory thalamus [82–84], in contrast with the spatial and temporally aligned modulation of thalamic activity by activation of CT afferents [78,85,86].

Despite these constraints, our study represents a significant advance toward understanding how intrathalamic connectivity shapes sensory processing and emergent thalamic dynamics. By providing a versatile and systematically constructed model, we offer a foundational framework that future experimental and computational efforts can refine and expand as new data become available.

### The role of intrathalamic connectivity in shaping angular tuning

Tuning maps are a fundamental aspect of sensory processing, extending far beyond the whisker system [87–96]. In nearly every sensory modality, neurons exhibit selective responses to specific stimulus features or ranges of features, emerging either at the single cell [27] or at the circuit level [93]. This neural selectivity allows the brain to represent, differentiate, and interpret a wide variety of external inputs, serving as a crucial building block for perceptual processes [97,98]. By examining tuning properties and maps across different sensory pathways, researchers gain insights into how neuronal populations encode complex stimuli and how these representations shape both local microcircuit dynamics and global network states involved in perception, attention, and behavior [99–101]. In the whisker pathway, angular tuning maps reveal how neurons transform and integrate sensory inputs, offering insights into the connectivity and activity patterns underlying information processing in response to whisker deflection [26,55].

In this study, we showed that topological inputs play a key role in shaping the angular tuning response of thalamic neurons in the whisker pathway. By incorporating experimentally observed topological projections from the brainstem to the thalamic barreloid [28] and systematically varying the intrathalamic connectivity, we were able to drive TC activity that reproduced angular tuning patterns measured in vivo [1] and showed that the angular tuning propagation from TC to TRN scales proportionally with the percentage of on-center feedforward TC→TRN connections.

Topological projections can enhance precision by creating localized, high-intensity firing zones for preferred stimulus features, thereby improving discrimination [1,26,27]. In contrast, less orderly projections (e.g., uniform connectivity) might yield more diffuse activity that could mask subtle differences in incoming signals, potentially reducing the specificity of the neuronal response [102]. By varying the model’s connectivity, we were able to shape the tuning curves to match the angular tuning observed experimentally [1]. While the connectivity from MLe→TC was set to match experimental data, we were able to shape the TC→TRN tuning by adjusting the percentage of on-center feedforward TC→TRN connections. This result is in line with previous studies indicating that the thalamic circuit in the whisker pathway is highly topological, showing a clustering of cells with similar angular preference along the depth of the thalamic barreloids [27,103]. Taken together, this suggests that angular tuning could be used as a biomarker to estimate the organization of TC→TRN connections.

Our model suggests that closed-loop TC↔TRN connectivity is better suited to support angular tuning, since introducing open-loop connections led to greater variability in the angular tuning maps (TRN→TC AT_RMSE_, Fig 5H). This impaired fidelity would be counterproductive in a system whose primary function during wakefulness is to relay information from the brainstem to the cortex.

### The role of TRN feedback in regulating sleep-state dynamics

The transition from wake to sleep is characterized by a global shift in circuit dynamics [7,48] and is driven by a series of neuromodulatory inputs that alter the ionic balance and conductance of transmembrane currents [9,38,40–42,62]. As the brain transitions into NREM sleep, thalamic neurons shift from the awake-associated tonic firing to burst firing [32,104]. They also show a decreased responsiveness to external stimuli, and the N2 stage of NREM is marked by the emergence of spindle oscillations [7,39,105].

Our model reveals that different connectivity configurations have a pronounced effect on spindles during sleep. We observed that increasing the percentage of open-loop connections results in a proportional decrease in mean LFP envelope amplitude, impairing the circuit’s ability to sustain robust spindle oscillations. In contrast, circuits with closed-loop and/or uniform connectivity preserve a higher spindle amplitude, showing that reciprocal closed-loop interactions between the TC and TRN neurons are supportive of spindle oscillations. This result aligns with physiological observations suggesting that thalamic circuits rely on coordinated feedback loops to promote synchronized firing and regulate sleep-related oscillations [8,32,106].

Additionally, our model showed coordinated activation of thalamic neurons during CT Up states, demonstrating that cortical inputs not only enhance thalamic excitability but also act as a timing signal that can lock the onset of spindle waves to particular phases of the slow oscillation [107].

In sum, these results suggest that while the whisker-related barreloid circuit may be structured to optimize angular tuning and sensory relay during wakefulness, these reciprocal connections can also be used to sustain an oscillatory mode during sleep. By allowing both robust spindle generation and proper inhibitory gating, the combined presence of uniform and closed-loop projections may offer a biologically advantageous balance: effective sensory relay when the system is awake and synchronization of sleep rhythms when neuromodulatory inputs shift the circuit into an oscillatory regime. This insight highlights the versatility of thalamic circuits and underscores the functional importance of TRN feedback in orchestrating the thalamo-cortical dynamics that underlie the sleep state.

### Corticothalamic influence in processing information

Coordinated CT feedback can significantly enhance the precision of sensory processing by selectively amplifying relevant inputs and suppressing irrelevant ones, thereby improving sensory discrimination [108,109]. It is also speculated that CT inputs can regulate the firing properties of TC neurons, selectively switching between tonic and burst firing [110]. Our simulations show that closed-loop CT connections enhance angular tuning by selectively boosting aligned neuronal responses. These results are in line with previous reports showing that neurons in the barrel cortex display a preferred direction, similar to the brainstem and thalamus [31,111], and reinforce the idea of a closed-loop CT connectivity.

Moreover, the different responses observed in the model’s TC and TRN populations suggest distinct roles of CT feedback across nuclei. The pronounced effect of CT modulation on TRN neurons highlights their critical role in gating and regulating sensory relay through selective inhibition [4,112]. In contrast, relay nuclei such as the TC neurons in VPM appear less directly modulated by CT input, likely reflecting dominant feedforward inputs (e.g., from brainstem) that inherently shape their responses [28,67].

As mentioned previously, we acknowledge that representing the CT feedback as a spiking network has its drawbacks compared with including a population of biophysically detailed L6 CT neurons. However, that only makes the model harder to constrain, and would only be reasonable in the context of including a full-scale model of the barrel column. Given that the main response of interest in our study is the angular tuning in the network, which is characterized within a 20 ms window following the whisker deflection, the effects of including a detailed model of the CT network would be significantly limited. A study from Hirai et al. shows the profile of this CT response with rich detail, describing the temporal dynamics of cortical neurons in a layer-specific fashion, highlighting how L6 CT neurons respond within the same temporal window of neurons in L4, and that this response is significantly limited to a time window of < 30 ms [45]. Therefore, we reasoned that a more direct strategy to test the effects of direct activation of L6 CT neurons would be to assume the transfer of angular tuned information from TC to CT neurons, which would in turn respond by propagating that activity back to thalamus.

Moreover, there are important circuit-level details within the L6 CT neurons that are not fully understood, with response profiles that are both state-dependent (varying between whisker-activated (16.5%), -suppressed (11.3%), or being silent (28.7%) or sparse (43.5%) firing) [71] and context-dependent [113,114]. We accounted for that variability by gradually increasing the number of angular-tuned CT neurons, showing that sharpening of angular tuning by direct CT activation was consistent across different levels of CT modulation.

These results collectively suggest a functional model in which selective CT activation serves to amplify sensory precision through targeted modulation of thalamic circuits. This idea has been proposed before [115], but this is the first demonstration of this mechanism in a thalamic circuit computational model. Moreover, the pronounced effect of CT modulation observed within the TRN supports a framework in which the TRN acts as a critical node for enhancing angular discrimination, potentially through selective inhibition of less relevant sensory signals [4,19,116]. This mechanism could facilitate improved signal-to-noise ratios and more precise sensory encoding in behavioral contexts. Future studies should explore the behavioral and perceptual consequences of such selective CT modulation.

### Investigating open- and closed-loop circuits through computational modeling

The first computational model of a thalamic relay neuron was proposed by McCormick and Huguenard [117], showing that thalamic dynamics could be reproduced *in silico*. Recent studies have modeled the dynamics of thalamic cells in greater detail, combining realistic morphology and biophysics of these neurons, as well as synaptic physiology and connectivity [52,54]. However, these studies focused on the principles of a general thalamo-cortical circuit, whose structure and synaptic organization differ significantly from the barreloid thalamus [118,119] and did not address the question of whether thalamic circuits are organized as open- or closed-loops, which remains unresolved [14,19].

Previous computational studies have addressed this question, demonstrating that open-loop circuits were suitable for sustaining the propagation of thalamo-cortical activity [69,70]. However, these studies used simplified circuit models composed of single-compartment neurons, optimized for signal propagation. In doing so, they did not consider the impacts of neuronal morphology [74] and the timing and distribution of presynaptic inputs [110,120] in the firing properties of the circuit.

In contrast, our highly detailed model captures realistic awake-state dynamics and physiologically accurate LFP signatures, providing insights into circuit-level phenomena such as dendritic integration, input segregation, and spatial coordination [52,121–123], which are not accessible in simplified circuit models. Moreover, our systematic investigation of intrathalamic connectivity using a detailed circuit shows that closed-loop connectivity can also support the propagation of oscillatory activity, previously attributed primarily to open-loop circuits (Fig 6H).

Identifying a general connectivity strategy across thalamic nuclei requires accounting for the unique characteristics of each sensory modality and their influence on network development and synaptic organization. In this context, open- and closed-loop circuits may coexist [19]: closed-loop circuits may support local processing of information and generation of sleep spindles, whereas open-loop motifs are responsible for facilitating long-range intrathalamic communication [124,125].

## Methods

This section describes model development with data provenance, major features of the final model, and the analysis and experimental methods. The full documentation of the final model and analyses is the source code itself.

### Data and code availability

The model source code, analysis scripts, and experimental data used for model constraints and validation are available on GitHub at: https://github.com/suny-downstate-medical-center/thalamus_netpyne. A permanent archive of the simulation output data generated in this study is available at https://doi.org/10.5281/zenodo.16888006. The model source code is fully compatible with the NetPyNE framework and can be exported to standardized formats such as SONATA and NeuroML.

Any additional information required to run or analyze the model is available from the **lead contacts** upon request.

### Thalamo-cortical circuit

The thalamo-cortical circuit reconstructed in this model includes: a thalamic barreloid composed of TC neurons representing the VPM nucleus; a section of the TRN composed of GABAergic neurons; and two whisker deflection models representing a brainstem barrelette and the CT neurons from an infrabarrel in L6A of S1 (Fig 1A). Circuit dimensions and number of cells were based on studies that reconstructed the anatomy and cell density of nuclei in the brainstem, thalamus, and cortex (Fig 1B) [25,68,126,127]. The resulting circuit model contains 1,560 cells, of which 562 are morphologically and biophysically detailed thalamic neurons (346 TC neurons, 216 TRN neurons), uniformly distributed across the volume of their respective nuclei (Fig 1B).

### Cell morphology and biophysics

We employed existing cell models from a state-of-the-art computational model of the non-barreloid thalamus simulated in NEURON [52]. To ensure morphological diversity, we sampled 5 different cell templates for each population (Fig 1C). We modeled the remaining neurons in the circuit as artificial spike-generators, representing the driver and modulatory excitatory inputs from a brainstem barrelette and a cortical infrabarrel, respectively (Fig 1B) [28,67,68].

We extracted the properties of the non-barreloid thalamic microcircuit using the *snap* Python package (v 0.13.0), available in the Blue Brain GitHub repository, and converted the cell models to the SWC format to import into NetPyNE.

In the original study, the cell models are tuned to a very low RMP and require a constant current-clamp input to set the membrane voltage to the desired level. With this in mind, we developed an algorithm to replicate the current-clamp input combined with an Ornstein-Uhlenbeck process, which are used to provide different current stimulations to set the RMPs across cells in the source model [52]. For that, we current-clamped each cell type with different subthreshold amplitudes without noise, and fitted an equation to map the current amplitude required to set the membrane at a given voltage (more details can be found in the **source code**). We validated the cell models using a current clamp protocol, testing the ability of the cells to generate tonic and burst firing under different conditions (Fig 1C).

### Model connectivity

Connectivity was defined based on studies that mapped well-known brainstem, intrathalamic, thalamo-cortical, and corticothalamic projections [14,15,17,18,27,28,43,67,118]. The parameters used to constrain and validate the circuit connectivity were extracted from experimental and modeling studies, and included the number of connections between cell types, the number of synapses per connection, and the spatial distribution of these synapses in the dendritic tree for each pathway [28,52,67,128–130]. The topology of these connections was also based on the pattern of innervation from axonal afferents projecting into each nucleus.

To match the number of projections to and from TRN observed experimentally, we added an external ring of TRN cells, representing the adjacent TRN sectors that project to a given thalamic barreloid [78]. Having this ring of TRN neurons also helps minimize the border effect, which is common in networks that use position-based connectivity rules. This ring comprises 106 (49%) of the 216 TRN cells driven by 10 Hz spiking activity to approximate the spiking of the biophysical TC neurons [52], and are excluded from the angular tuning and spindle calculations.

The model also features gap junction connectivity between TRN neurons based on overlapping dendritic compartments. We devised an algorithm to detect the distance between neuronal compartments to create the pairwise connections based on a maximum distance threshold. Following, to match experimental observations, we adjusted the distance threshold (set to 5 µm) and synaptic pruning (maximum of 20 gap junction connections per cell). We validated the model’s gap junction distribution profile against experimental data [52,53] (Fig 1F).

In the case of TC↔TRN connectivity, studies are inconclusive about the arrangement of their axonal projections [19]. Therefore, we studied different intrathalamic connectivity configurations in this pathway to evaluate its influence on angular tuning responses and sleep dynamics. We started with a uniform connectivity between TC↔TRN (Fig 2), and later compared against the other circuit configurations (Figs 5-7).

The resulting circuit model comprised approximately 690k synapses, distributed over 95k cell-to-cell connections. Specifically, the uniform circuit had 95,558 connections and 691,162 synapses; the open-loop circuit, 91,939 connections and 690,450 synapses; the closed-loop circuit, 95,741 connections and 691,363 synapses; and all circuits share the same 3,432 gap junctions. Fig 1 compares the connectivity of the current model and a non-barreloid thalamic model [52] based on the ratio of projections between different pathways (Fig 1E) and the distribution of gap junctions (Fig 1F).

### Synaptic models

All chemical synapses included short-term plasticity implemented using the Tsodyks-Markram model [76]. The values of synaptic conductance and short-term plasticity (Use, Depression, and Facilitation) for each pathway were sampled from a truncated normal distribution following the parameters of the source model (Fig 1D) [52]. The weights of these connections were tuned within physiological ranges to obtain different circuit dynamics. Each gap junction was implemented as a 0.2 nS bidirectional conductance between the two compartments for all connections, following the source implementation (Fig 1F) [52].

### Brainstem whisker deflection model

We used the response of brainstem neurons during passive whisker deflection to develop a model that simulates realistic spiking activity from a brainstem barrelette encoding whisker deflection inputs to the thalamus [2]. This brainstem model allows the selection of different deflection angles, intervals, and total duration, as well as the number of input sources representing each brainstem neuron. The brainstem model used in this study consists of 180 artificial neurons, modeled as spike generators, each representing a 2° sector in polar coordinates [27,66]. The spiking activity of these neurons is derived from PSTHs recorded in the brainstem and thalamus (Fig 3A), encoding angular-tuned information by scaling each neuron’s response based on the angle of whisker deflection (Fig 3B) [2,27].

The PSTH quantifies the spike probability for a given group of cells over time (Fig 3A). During baseline, the spiking activity was modeled based on the mean and standard deviation of the PSTH response. We fitted the ON (deflection onset) and OFF (release) transition periods of the deflection events to the PSTH data using triple and double exponential decay functions, respectively (Fig S3A). The angular tuning response quantifies the direction selectivity of the cells in the circuit (Fig 3B). Experimentally, it is obtained by mapping the preferred direction of each recorded cell, which is used as a reference angle, followed by measuring the overall response of all cells to deflections in all different directions [2,27]. To replicate that tuning in our model, each cell is assigned a preferred angle of maximal responsiveness (0°–358°), based on its relative position along the y-axis. During baseline, all cells have a similar firing rate, extracted from the PSTH data. During a deflection event, this firing probability is scaled based on the relative distance between the cell’s preferred angle and the deflection angle. Lastly, the probability is rescaled to match the area under the curve before angular tuning scaling (Fig S3B). This rescaling preserves the angular tuning of the cells while reproducing the overall firing rate of the population.

This implementation is demonstrated in Fig 3C and summarized by the following equation:

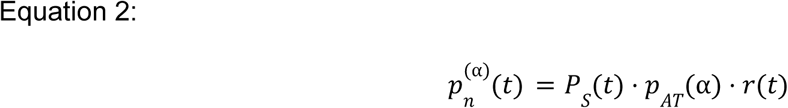

Where:

*p*_*n*_^(α)^(*t*) is the final spike probability for a given cell *N*, which encodes the angle *n*, during a whisker deflection towards angle α; *P*_*S*_ (*t*) is the population spike probability, the same for all cells and time-dependent (extracted from PSTH data); *p*_*AT*_ (α) is the angular tuning probability scaling factor, based on the distance ∣*n* − α∣; *r*(*t*) is a rescaling factor that adjusts the area under the spike probability curve after tuning.

The values of *p*_*AT*_(*t*) and *r*(*t*) are defined as follows:

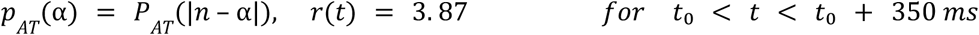

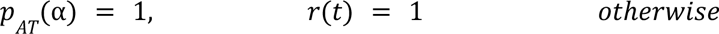

Where:

*P*_*AT*_ (|*n* – α|) is the function that determines the value of *p*_*AT*_ (α), which is a scaling factor used to modify the spike probability based on the distance from the neuron that encodes the angle *n* to the target deflection angle α.

These equations are used to create a dataset for the 180 cells, combining the PSTH and the angular tuning responses. For each deflection event, the probability of firing changes from the baseline value to the ON and OFF values, which are scaled based on the distance between the angle encoded by a given cell and the target deflection angle. During each run of a 16-second simulation (Fig S3C), the thalamic circuit received 14 deflection events of 250 ms duration for the same angle. We ran the simulations for 8 angles, starting at 0° and spaced by 45° (Fig 3D-E). We calculated the angular tuning response for each population by quantifying the spike output of each cell during a 20-ms window based on their position along the vertical axis (Fig S3C). The spikes were grouped based on the preferred direction of each group of cells, and the spiking output of each direction was normalized by the spike output of the target angle (Fig S3C). Lastly, we aligned the normalized data in the 270° direction to create a combined angular tuning map (Fig S3D). We verified the model’s fit against the experimental data using a Wilcoxon signed-rank test, comparing each value along the angular tuning map for the experimental data with the distribution of values for the same angle after aligning to a common direction. We found no statistical difference in 5 out of 8 directions (0°, 45°, 225°, 270°, and 315°), which included the angles that were aligned with the direction of the deflection (Table S5i). Additionally, the difference in distribution of spikes between the non-significant and significant directions was only 3.47%, showing that the model provides a good fit to the dataset (Table S5ii). The result was a realistic stimulation dataset that mimicked the topological information encoded in the brainstem and was used to drive the activity in the thalamic populations.

### Corticothalamic feedback

The CT feedback in the model was implemented as 818 artificial neurons, or spike generators, firing at 5 Hz, consistent with experimental estimates [52,68,71], and was used to characterize CT feedback effects on thalamic sleep/wake and angular tuning responses.

In the awake state, CT inputs consisted of 5 Hz activity in the spike generator population. For the CT Up/Down modulation in the sleep state, the spike generators alternated between 5 Hz (500 ms) and 1 Hz (1500 ms). For the angular-tuned CT modulation simulation, either 0%, 10%, 25%, 50%, or 100% of the 5 Hz spike generators were replaced with angular-tuned spike generators and evenly distributed across the CT population. Angular tuning in CT neurons was implemented using the brainstem whisker deflection model.

### Circuit-level modifications and model tuning

Because this model is a reimplementation of an already tuned circuit, we expected that as we rescaled the size of the circuit, the strength of some synaptic pathways would require retuning. After an initial exploration in the parameter space of RMP values, we set the TC neurons to -61 mV and the TRN neurons to -71 mV using the current-clamping algorithm and added the Ornstein-Uhlenbeck noise [52]. With the circuit connected, we modified the conductance values for all pathways using a ‘weight’ variable that scales the postsynaptic potential of the selected pathways, exploring a range of values between 0.1 and 5.0. We used the firing rate of the TC and TRN neurons to establish a fixed value for the AMPA/NMDA strength for the different pathways across sleep and wake states. Our goal was to reproduce the oscillatory activity in the spindle frequency range characteristic of thalamo-cortical circuits in sleep. Following that, we applied the same strategy to reproduce the wake state dynamics, depolarizing the TC and TRN RMPs (-58 mV) and rescaling their ionic currents and GABA strength, following literature reports [9,38,40–42,62]. All changes are summarized in Table 1.

### Model implementation and simulation

This model was developed, simulated, optimized and analyzed using Python (version 3.9.7) and the NetPyNE tool (version 1.0.6) [131], an open-source framework that streamlines the modeling of large-scale biophysical neural circuits, and which uses the NEURON simulator (version 7.8.2) as simulation engine [132,133].

Simulations were executed on a local supercomputer at SUNY Downstate, and each full run of the model required 100.62 hours to complete, running on 8 machines with 64 cores each (32,320 core hours). In these conditions, each instance of the model, consisting of a 16-second simulation, required approximately 2h35’ to complete, and demanded approximately 128 GB of RAM.

### Key resources table

| RESOURCE | SOURCE | IDENTIFIER |
| --- | --- | --- |
| **Deposited data** |  |  |
| Thalamo-cortical model output simulated data | https://doi.org/10.5281/zenodo.16888006 |  |
| **Software and algorithms** |  |  |
| Thalamo-cortical model and data analysis source code | https://github.com/suny-downstate-medical-center/thalamus_netpyne;<br>This paper |  |
| NetPyNE | http://netpyne.org | RRID: SCR_014758 |
| NEURON | https://neuron.yale.edu | RRID: SCR_017449 |
| snap | https://github.com/BlueBrain/snap | N/A |
| Python | http://python.org | RRID: SCR_008394 |
| SciPy | https://scipy.org | RRID: SCR_008058 |

## Acknowledgements

This work was supported by the National Institutes of Health under award numbers NIH U24EB028998, NIMH P50MH109429, NYS SCIRB DOH01-C38328GG, all to S.D., and by the National Institute of Mental Health under Award Number R56MH132637 to C.V.

## Supporting information

Supporting information can be found online along with the manuscript.

## Author contributions

Conceptualization, J.V.S.M., S.D.-B.; Data curation, J.V.S.M., F.S.B., Z.A., S.D.-B.; Formal analysis, J.V.S.M., F.S.B., Z.A., S.D.-B.; Funding acquisition, S.D.-B.; Investigation, J.V.S.M., F.S.B., Z.A., S.R.C., C.V., S.D.-B.; Methodology, J.V.S.M., F.S.B., Z.A., S.R.C., C.V., S.D.-B.; Project administration, S.D.-B.; Resources, J.V.S.M., S.D.-B.; Software, J.V.S.M., F.S.B., Z.A.; Supervision, S.R.C., C.V., S.D.-B.; Validation, J.V.S.M., F.S.B., Z.A., S.R.C., C.V., S.D.-B.; Visualization, J.V.S.M.; Writing - Original Draft Preparation, J.V.S.M.; Writing - Review & Editing, J.V.S.M., F.S.B., Z.A., S.R.C., C.V., S.D.-B.

---

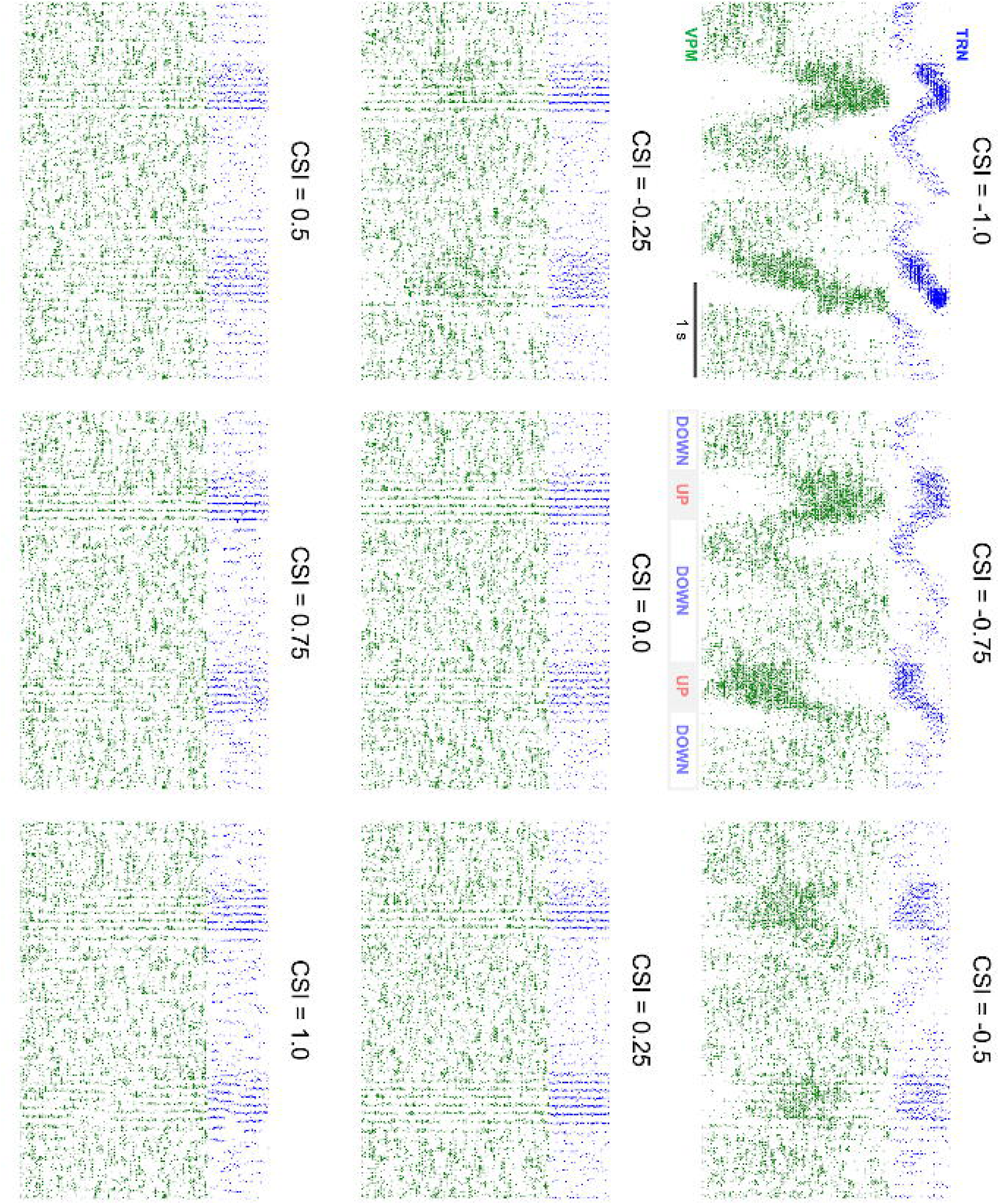

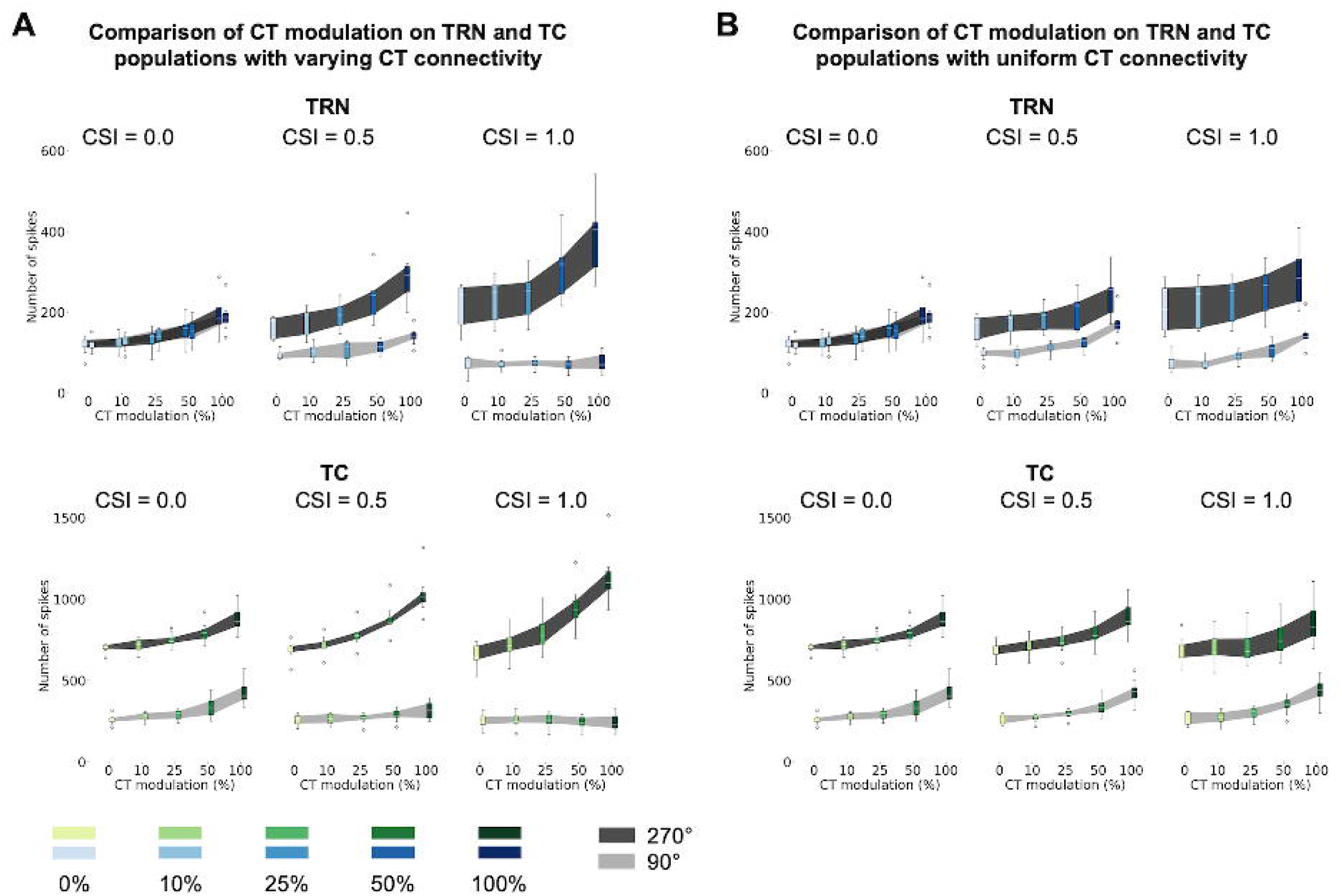

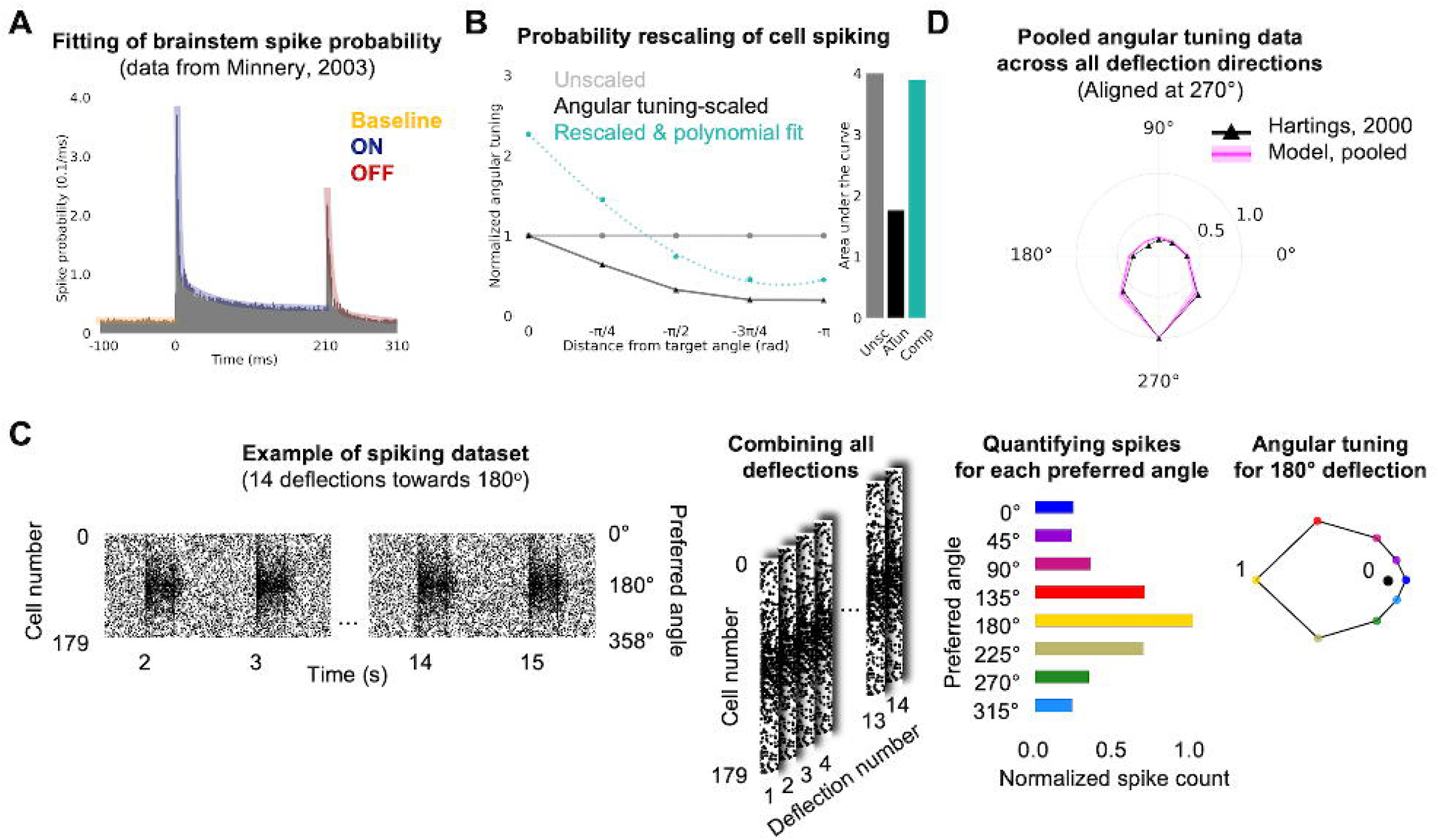

