## Supporting information for "Closed-Loop Connectivity Best Supports Angular Tuning and Sleep Dynamics in a Biophysical Thalamocortical Circuit Model"

### Statistical analysis

Tables S1, S2, S3, and S4 include details of the statistical analysis for Figs 5G-H, 6H, 7H-I and 8E-F presented in the *Results*. All tests were performed in Python (version 3.9.7) using the SciPy package (version 1.10.1).

Table S5 includes the details of the statistical tests on the angular tuning of the whisker deflection spiking model.

**Table S1: Statistical tests for the effect of intrathalamic connectivity in the angular tuning**

***i)* Spiking activity of aligned (270°) vs opposing cells (90°) for different CSI values (Fig 5G)**

| **Pop** | *-1.0* | *-0.75* | *-0.5* | *-0.25* | *0.0* | *0.25* | *0.5* | *0.75* | *1.0* |
|---|---|---|---|---|---|---|---|---|---|
| *TRN* | 0.0009*** | 0.0009*** | 0.0009*** | 0.0030** | 0.95 | 0.0237* | 0.0009*** | 0.0009*** | 0.0001*** |
| *TC* | 0.0001*** | 0.0001*** | 0.0009*** | 0.0009*** | 0.0009*** | 0.0001*** | 0.0001*** | 0.0009*** | 0.0001*** |

***ii)* Root mean squared error of CSI tested vs CSI = 0.0 (Fig 5H)**

| **Pop** | *-1.0* | *-0.75* | *-0.5* | *-0.25* | *0.25* | *0.5* | *0.75* | *1.0* |
|---|---|---|---|---|---|---|---|---|
| *TRN* | 0.0006*** | 0.0006*** | 0.0006*** | 0.0006*** | 0.0006*** | 0.0006*** | 0.0006*** | 0.0006*** |
| *TC* | 0.0006*** | 0.0006*** | 0.0006*** | 0.0006*** | 0.6200 | 0.1649 | 0.1282 | 0.0379* |

***iii)* Root mean squared error of CSI opposite values**

| **Pop** | *-0.25 vs 0.25* | *-0.5 vs 0.55* | *-0.75 vs 0.75* | *-1.0 vs 1.0* |
|---|---|---|---|---|
| *TRN* | 0.1282 | 0.5347 | 0.3176 | 0.2593 |
| *TC* | 0.0006*** | 0.0006*** | 0.0006*** | 0.0023** |

Two-sided Mann-Whitney U test
p >= 0.05: ns ; p < 0.05: * ; p < 0.01: ** ; p < 0.001: *** ; p < 0.0001: ****

**Table S2: Statistical tests for the effects of TRN and TC mean LFP amplitude in Up and Down states across different circuits - Fig 6H**

***i)* CSI tested vs CSI = 0.0**

| State | Pop | *-1.0* | *-0.75* | *-0.5* | *-0.25* | *0.25* | *0.5* | *0.75* | *1.0* |
|---|---|---|---|---|---|---|---|---|---|
| ***Up*** | *TRN* | 0.0006*** | 0.0006*** | 0.0041** | 1.0 | 0.9015 | 0.4557 | 0.6200 | 0.8048 |
| | *TC* | 0.0006*** | 0.0006*** | 0.0006*** | 0.1282 | 1.0 | 1.0 | 0.2086 | 0.1650 |
| ***Down*** | *TRN* | 0.9015 | 0.2593 | 0.4557 | 0.8048 | 0.8048 | 0.8048 | 0.5350 | 0.0262* |
| | *TC* | 0.2593 | 0.3829 | 1.0 | 0.9015 | 0.6200 | 0.3827 | 0.9015 | 0.1282 |

***ii)* CSI opposite values**

| State | Pop | *-0.25 vs 0.25* | *-0.5 vs 0.55* | *-0.75 vs 0.75* | *-1.0 vs 1.0* |
|---|---|---|---|---|---|
| ***Up*** | *TRN* | 0.9015 | 0.1282 | 0.0006*** | 0.0023** |
| | *TC* | 0.0973 | 0.0006*** | 0.0006*** | 0.0006*** |

***iii)* Up vs Down states for different CSI values**

| Pop | *-1.0* | *-0.75* | *-0.5* | *-0.25* | *0.0* | *0.25* | *0.5* | *0.75* | *1.0* |
|---|---|---|---|---|---|---|---|---|---|
| *TRN* | 0.3829 | 0.0262* | 0.0006*** | 0.0006*** | 0.0006*** | 0.0006*** | 0.0006*** | 0.0006*** | 0.0262* |
| *TC* | 0.0070** | 0.0262* | 0.1282 | 0.0728 | 0.0006*** | 0.0006*** | 0.0006*** | 0.0006*** | 0.0023** |

Two-sided Mann-Whitney U test
p >= 0.05: ns ; p < 0.05: * ; p < 0.01: ** ; p < 0.001: *** ; p < 0.0001: ****

**Table S3: Statistical tests for changes in spiking activity of TC and TRN populations under closed-loop CT modulation - Fig S2A**

***i)* Spiking activity of aligned (270°) vs opposing cells (90°) for different levels of CT modulation**

| Pop | CSI | *0%* | *10%* | *25%* | *50%* | *100%* |
|---|---|---|---|---|---|---|
| *TRN* | *0.0* | 0.9581 | 0.9581 | 0.7927 | 0.8746 | 0.9581 |
| | *0.5* | 0.0009*** | 0.0003*** | 0.0002*** | 0.0009*** | 0.0009*** |
| | *1.0* | 0.0009*** | 0.0009*** | 0.0009*** | 0.0002*** | 0.0002*** |
| *TC* | *0.0* | 0.0009*** | 0.0002*** | 0.0002*** | 0.0002*** | 0.0002*** |
| | *0.5* | 0.0002*** | 0.0002*** | 0.0009*** | 0.0002*** | 0.0002*** |
| | *1.0* | 0.0002*** | 0.0002*** | 0.0009*** | 0.0002*** | 0.0002*** |

***ii)* Spiking activity of CT tested vs CT = 0.0%**
**for aligned and opposite neurons of different circuit configurations**

| Pop | Angle | CSI | *10%* | *25%* | *50%* | *100%* |
|---|---|---|---|---|---|---|
| *TRN* | *Aligned* | *0.0* | 0.6454 | 0.1412 | 0.0281* | 0.0027** |
| | | *0.5* | 0.1605 | 0.0830 | 0.0054** | 0.0002*** |
| | | *1.0* | 0.5280 | 0.4306 | 0.0404* | 0.0019** |
| | *Opposite* | *0.0* | 0.4616 | 0.0656 | 0.0379* | 0.0003*** |
| | | *0.5* | 0.3181 | 0.2698 | 0.0308* | 0.0016** |
| | | *1.0* | 0.8333 | 0.9580 | 0.6363 | 0.9162 |
| *TC* | *Aligned* | *0.0* | 0.5737 | 0.0047** | 0.0011** | 0.0002*** |
| | | *0.5* | 0.1304 | 0.0156* | 0.0003*** | 0.0002*** |
| | | *1.0* | 0.1304 | 0.0104* | 0.0002*** | 0.0002*** |
| | *Opposite* | *0.0* | 0.2698 | 0.1028 | 0.0238* | 0.0009*** |
| | | *0.5* | 0.9162 | 0.8335 | 0.1605 | 0.0830 |
| | | *1.0* | 0.6454 | 0.7927 | 0.7525 | 0.6454 |

Two-sided Mann-Whitney U test
p >= 0.05: ns ; p < 0.05: * ; p < 0.01: ** ; p < 0.001: *** ; p < 0.0001: ****

**Table S4 - Statistical tests for changes in spiking activity of TC and TRN populations under uniform CT modulation - Fig S2BF**

***i)*** **Spiking activity of aligned (270°) vs opposing cells (90°) for different levels of CT modulation**

| **Pop** | **CSI** | *0%* | *10%* | *25%* | *50%* | *100%* |
|---|---|---|---|---|---|---|
| *TRN* | *0.0* | 0.9581 | 0.9581 | 0.7927 | 0.8746 | 0.9581 |
| | *0.5* | 0.0009*** | 0.0009*** | 0.0009*** | 0.0027** | 0.0047** |
| | *1.0* | 0.0002*** | 0.0009*** | 0.0002*** | 0.0002*** | 0.0019** |
| *TC* | *0.0* | 0.0009*** | 0.0002*** | 0.0002*** | 0.0002*** | 0.0002*** |
| | *0.5* | 0.0002*** | 0.0002*** | 0.0002*** | 0.0002*** | 0.0002*** |
| | *1.0* | 0.0002*** | 0.0009*** | 0.0002*** | 0.0009*** | 0.0002*** |

***ii)*** **Spiking activity of CT tested vs CT = 0.0% for aligned and opposite neurons of different circuit configurations**

| **Pop** | **Angle** | **CSI** | *10%* | *25%* | *50%* | *100%* |
|---|---|---|---|---|---|---|
| *TRN* | *Aligned* | *0.0* | 0.6454 | 0.1412 | 0.0281* | 0.0027** |
| | | *0.5* | 0.5054 | 0.1412 | 0.8290 | 0.0047** |
| | | *1.0* | 0.5737 | 0.4418 | 0.1149 | 0.0519 |
| | *Opposite* | *0.0* | 0.4616 | 0.0656 | 0.0379* | 0.0003*** |
| | | *0.5* | 0.7921 | 0.0582 | 0.0135* | 0.0009*** |
| | | *1.0* | 0.7126 | 0.0927 | 0.0207* | 0.0013** |
| *TC* | *Aligned* | *0.0* | 0.5737 | 0.0047** | 0.0011** | 0.0002*** |
| | | *0.5* | 0.2786 | 0.1716 | 0.0207* | 0.0006*** |
| | | *1.0* | 0.5632 | 0.7209 | 0.1560 | 0.0047** |
| | *Opposite* | *0.0* | 0.2698 | 0.1028 | 0.0238* | 0.0009*** |
| | | *0.5* | 0.6742 | 0.0830 | 0.0063** | 0.0001*** |
| | | *1.0* | 0.5632 | 0.1149 | 0.0100* | 0.0011** |

Two-sided Mann-Whitney U test
p >= 0.05: ns ; p < 0.05: * ; p < 0.01: ** ; p < 0.001: *** ; p < 0.0001: ****

**Table S5: Experimental data vs whisker deflection model**

***i)* Wilcoxon signed-rank test**

| **Angle** | 0 | 45 | 90 | 135 | 180 | 225 | 270 | 315 |
|---|---|---|---|---|---|---|---|---|
| **Significance** | 0.7422 | 0.4609 | 0.0234* | 0.0078** | 0.0078** | 0.1094 | - | 0.1953 |

Wilcoxon signed-rank test: $p >= 0.05$: ns ; $p < 0.05$: * ; $p < 0.01$: **

***ii)* Difference in the number of spikes**

| | **Percentage of spikes** | | **Spike difference** |
|---|---|---|---|
| | ***Experimental*** | ***Model*** | ***Exp - Mod*** |
| **Not-significant angles** (0°, 45°, 225°, 270°, and 315°) | 72.95% | 69.48% | -3.47% |
| **Significant angles** (90°, 135°, 180°) | 27.05% | 30.52% | 3.47% |

### Supplementary figures

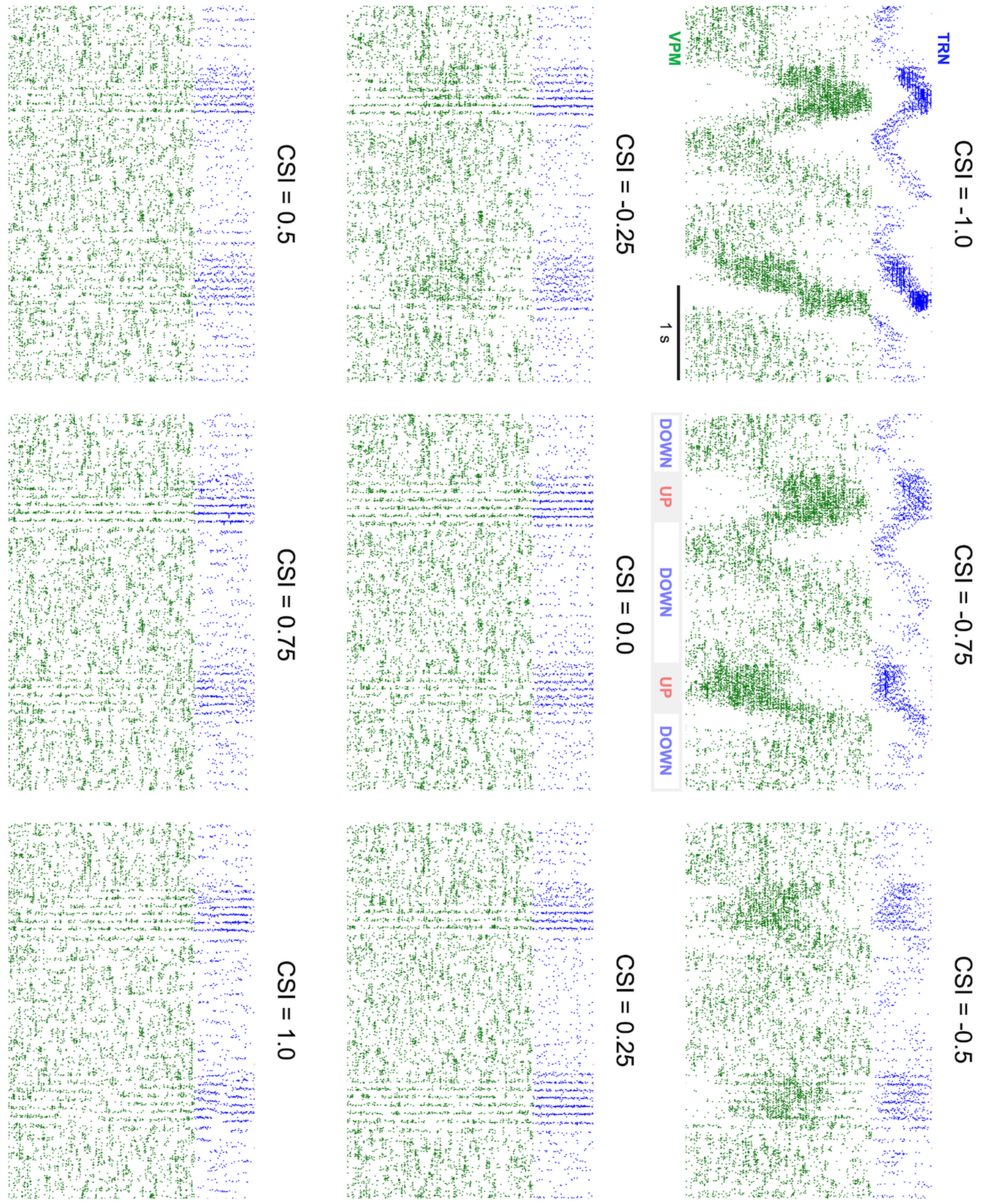


**Fig S1: Effect of intrathalamic connectivity (TC↔TRN) in the propagation of thalamocortical activity during Up and Down states.** A higher proportion of open-loop connections supports propagation of activity, in a traveling wave-like fashion, while uniform and closed-loop circuits promote network synchronization, sustaining spindle-like activity.

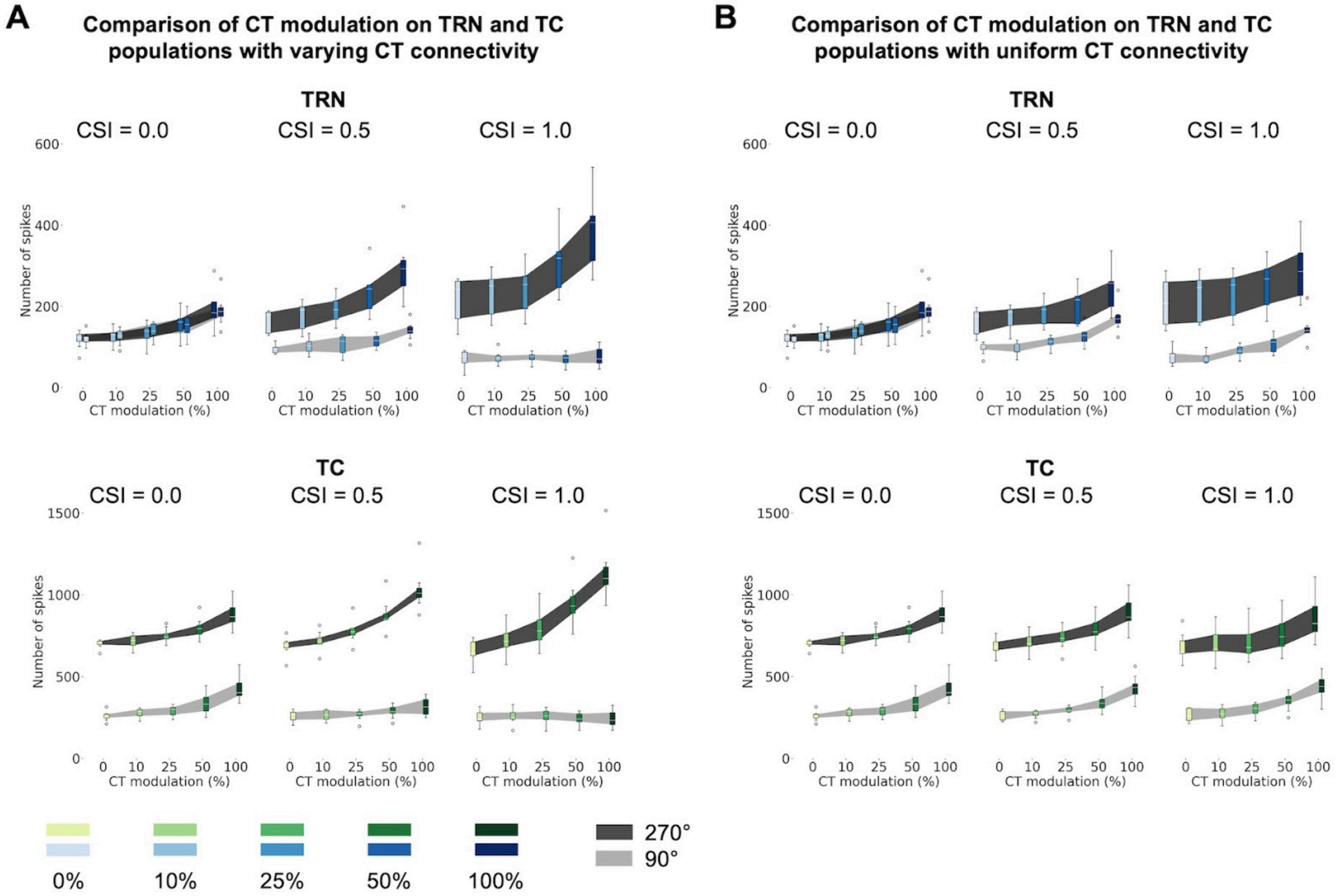


**Fig S2: Comparison on the effect of CT modulation in the angular tuning of TRN and TC populations.**

Comparison between the spiking of aligned and opposite neurons for TRN (A) and TC (B). See Tables S3 and S4 in *Supporting information* for detailed statistical analyses. TC: thalamocortical; TRN: thalamic reticular nucleus; CT: corticothalamic; CSI: connectivity symmetry index.

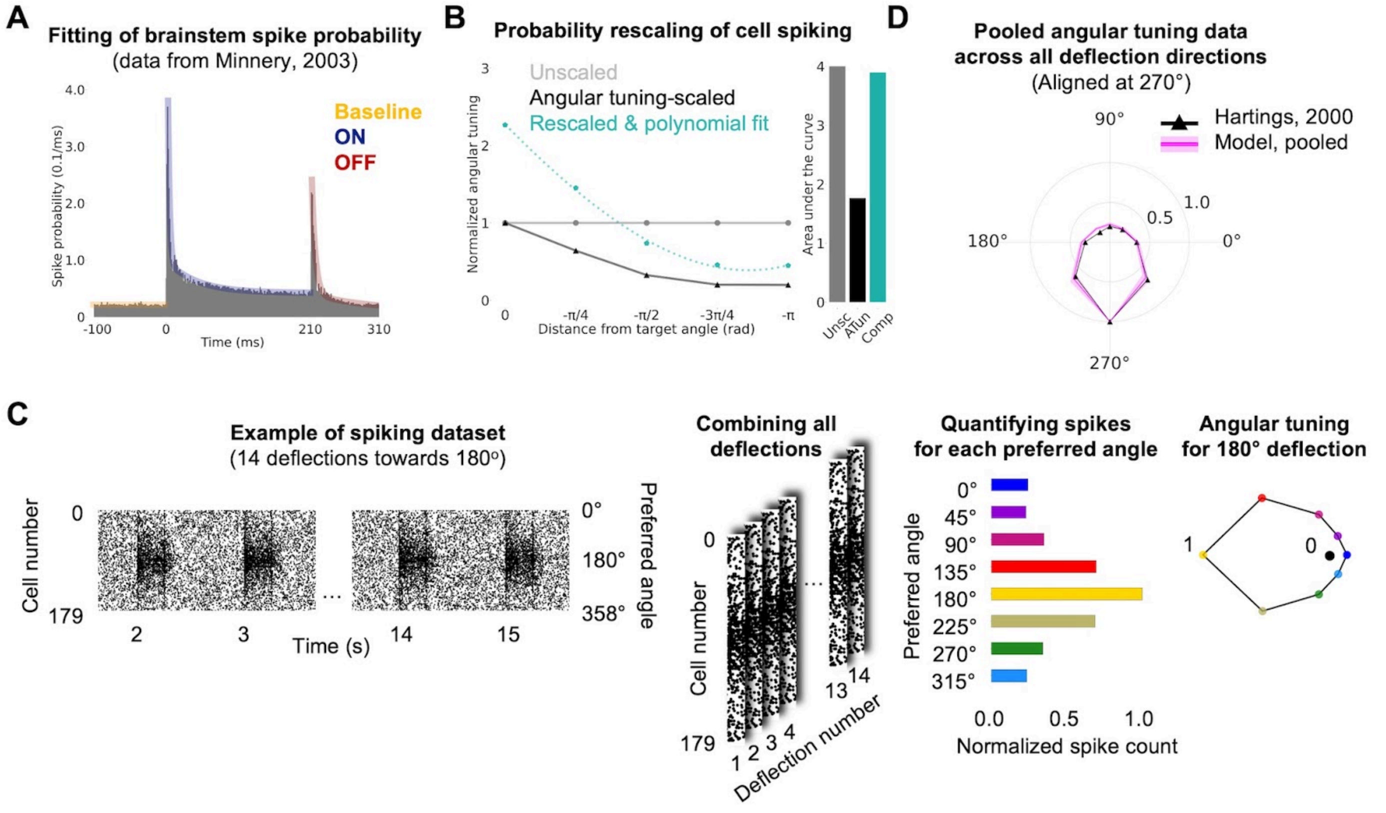


**Fig S3: Detailed description of the angular tuning calculation to validate the brainstem whisker deflection model**

(A–B) Fitting of experimental brainstem activity data during whisker deflection, including (A) spike probability (PSTH data from Minnery et al., 2003) and (B) angular tuning (Hartings et al., 2000). (C) Protocol for generating spiking datasets representing whisker deflections at eight angles (0° to 315° in 45° increments). We selected the first 20 ms of spiking activity for each deflection event to calculate the angular tuning. The spikes are combined and grouped based on the closest of the eight deflection angles, and for each angle, the total number of spikes is normalized based on the direction of the whisker deflection (set to 1.0). (D) Average angular tuning map derived from the whisker deflection model compared to experimental data (aligned to 270° reference angle from Hartings et al., 2000).
